# The hepatitis E virus capsid protein ORF2 counteracts cell-intrinsic antiviral responses to enable persistence in hepatocytes

**DOI:** 10.1101/2025.02.03.636239

**Authors:** Ann-Kathrin Mehnert, Sebastian Stegmaier, Carlos Ramirez, Vladimir Gonçalves Magalhães, Carla Siebenkotten, Jungen Hu, Ana Luisa Costa, Daniel Kirrmaier, Michael Knop, Xianfang Wu, Thibault Tubiana, Carl Herrmann, Marco Binder, Viet Loan Dao Thi

## Abstract

Hepatitis E virus (HEV) is a significant human pathogen causing both acute and chronic infections worldwide. The cell-intrinsic antiviral response serves as the initial defense against viruses and has been shown to be activated upon HEV infection. HEV can replicate in the presence of this response, but the underlying mechanisms remain poorly understood.

Here, we investigated the role of the HEV structural proteins ORF2 and ORF3 in immunocompetent cells. Mechanistically, we validated that ectopic ORF2, but not ORF3, interfered with antiviral and inflammatory signaling downstream of pattern recognition receptors, in part through interaction with the central adaptor protein TANK binding kinase 1. In the full-length viral context, ORF2 contributed to a reduced antiviral response and consequently, more efficient viral replication. In addition, we discovered a protective mechanism mediated by ORF2 that shielded viral replication from antiviral effectors. Using single-cell RNA-sequencing, we confirmed that the presence of ORF2 in infected cells dampened antiviral responses in both actively infected cells and bystanders. As a consequence, we found that early in the infection process, the progression of authentic HEV infection relied on the presence of ORF2, facilitating a balance between viral replication and the antiviral response within immunocompetent cells. Altogether, our findings shed new light on the multifaceted role of ORF2 in the HEV life cycle and improve our understanding of the determinants that may contribute to HEV persistence.

**Significance statement:** Hepatitis E virus (HEV) is an important yet often underestimated pathogen. Depending on the genotype, HEV infections can progress to chronicity, but the underlying mechanisms remain poorly understood. To gain insight into potential determinants, we investigated how HEV evades the body’s first line of defense, the cell-intrinsic antiviral response. We discovered that the HEV capsid protein ORF2 is crucial in limiting this response by interfering with antiviral signaling pathways and shielding viral replication from immune effectors. This balance between viral replication and the antiviral response contributes to persistent HEV infection in immunocompetent cells. Our findings reveal a new role for the HEV capsid protein in the viral life cycle and highlight it as an important target for novel therapeutic approaches.

## Introduction

Hepatitis E virus (HEV) is one of the major causes of acute hepatitis across the globe, affecting an estimated 20 million people every year (reviewed in^1,2^). As part of the *Hepeviridae* family, the *Paslahepevirus balayani* genus comprises eight genotypes of which HEV-1 to HEV-4, and more recently HEV-7, have been associated with human infections. HEV-1 and HEV-2 are transmitted fecal-orally and are restricted to humans. In contrast, HEV-3 and HEV-4 spread to humans zoonotically through consumption of meat products from domestic pigs or wild boar. Although HEV-1/-2 infections are usually self-limiting, they are associated with high mortality in pregnant women. Further, HEV-3 and -4 infection of immunocompromised individuals can result in chronicity and development of liver fibrosis and cirrhosis.

HEV is ingested and excreted as a naked virus but circulates in the bloodstream wrapped in a host membrane-derived quasi-envelope, which is acquired during budding from cells (reviewed in^3^). It has a positive-sense, single-stranded RNA genome of approximately 7.2 kilobases, encompassing three open reading frames (ORFs) that give rise to three viral proteins (reviewed in^2^). ORF1 contains the domains involved in viral replication, such as the RNA-dependent RNA polymerase (RdRp). ORF3 is a small phosphoprotein that mediates secretion of viral progeny.

Three distinct isoforms of ORF2 have been described in HEV-infected cells^4–7^. Owing to its N-terminal signal peptide, ORF2 can be secreted along the secretory pathway where it gives rise to the glycosylated (ORF2g) and cleaved (ORF2c) isoforms. ORF2g is secreted as a dimer^5,6^ and serves as an immunological decoy *in vivo*^5^. It has been postulated that positively charged residues within an arginine-rich motif (ARM; five arginine residues: RRRGRR) downstream of the signal peptide assist ORF2 in retaining a cytosolic orientation at the endoplasmic reticulum membrane^4^. The intracellular ORF2 isoform (ORF2i) assembles into infectious progenies. ORF2i is mainly located in the cytosol but can translocate to the nucleus, and is likely involved in many virus-host interactions^4,7^. Another study highlighted the importance of two ORF2 start codons. The first start codon can give rise to the secreted ORF2g, while the second gives rise to ORF2i with intracellular localization^5^. Although it remains unclear whether differential usage of these start codons is relevant in HEV infection, they can be used to abrogate expression of individual or all ORF2 isoforms.

Cell-intrinsic defense strategies are initiated by recognition of pathogen-associated molecular patterns (PAMPs) through pattern recognition receptors (PRRs) (reviewed in^8^). In epithelial cells, double-stranded (ds)RNA, the replication intermediate of RNA viruses, is detected by the family of retinoic acid-inducible gene I (RIG-I)-like receptors (RLRs) in the cytosol, including RIG-I and melanoma differentiation-associated protein 5 (MDA5), which are additionally regulated by laboratory of genetics and physiology 2 (LGP2). In the endosome, Toll-like receptor 3 (TLR3) recognizes dsRNA. Different adaptor proteins mediate downstream signaling: Toll/interleukin-1 receptor (TIR) domain-containing adaptor-inducing interferon-β (TRIF) is recruited by TLR3, while RIG-I and MDA5 induce polymerization of mitochondrial antiviral signaling protein (MAVS). Both result in activation of TANK binding kinase 1 (TBK1) and the inhibitor of nuclear factor-κB (IκB) kinase (IKK) complex. TBK1 phosphorylates interferon regulatory factor 3 (IRF3), which translocates into the nucleus and induces expression of type I and type III interferons (IFNs) and IFN-stimulated genes (ISGs). Autocrine and paracrine IFN signaling through Janus kinase/signal transducers and activators of transcription (JAK/STAT) result in expression of hundreds of ISGs with antiviral functions (reviewed in^9^). Activation of the IKK complex leads to nuclear translocation of nuclear factor-κB (NF-κB) and expression of inflammatory cytokines.

Despite growing interest and advances in recent years, the cell-intrinsic antiviral response to HEV infection remains poorly described. HEV-infected chimpanzees had a similar but weaker ISG response compared to hepatitis C virus (HCV) infection, likely owing to the lower HEV viremia^10^. Infection of innate immunocompetent hepatocyte models with HEV induced robust type III IFN responses, which were dependent on viral replication^11,12^. The induced IFN response appeared to be persistent but was unable to eliminate viral replication within the observed time frames^11,12^. Blunting of IFN induction did not result in enhanced viral replication, suggesting that HEV has developed mechanisms to persistently replicate in the presence of a sustained antiviral response^12^. Corroborating this, several studies have shown that HEV replication is relatively resistant to exogenous IFN treatment, as compared to, for example, HCV^11,13–15^. Yin and colleagues proposed that this is the result of an internal IFN refractoriness due to persistent activation of JAK/STAT1 signaling and retention of phosphorylated STAT1 in the cytosol rather than a direct viral antagonism^11^.

Nonetheless, all HEV proteins have previously been suggested to antagonize the cell-intrinsic antiviral response (reviewed in^16^). Within ORF1, the X domain, the putative papain-like cysteine protease (PCP) domain, and the methyltransferase (MeT) domain were shown to interfere with type I IFN induction and NF-κB-mediated signaling^17–20^. A combined MeT-Y-PCP polyprotein was further found to interfere with the JAK/STAT pathway^21^. The ORF2 protein has been reported to interact with TBK1^22,23^ and hinder activation of NF-κB^4,24^. Studies of potential ORF3-mediated antagonisms have been controversial: while some studies suggested that ORF3 interferes with type I IFN production, NF-κB-, and IFN-mediated signaling^25–27^, one study showed that ORF3 could enhance IFN induction^28^. However, most investigations have relied on overexpression of individual HEV proteins or subdomains of ORF1, and the physiological relevance of each antagonism for HEV replication remains elusive.

In the present study, we sought to clarify the role of HEV ORF2 and ORF3 proteins in antagonizing antiviral responses and their contribution to the progression of HEV infection. To address this, we made use of viral mutants and assessed differences in IFN and ISG responses upon authentic infection of different immunocompetent hepatocellular systems in bulk and at single-cell resolution. We identified a replication-limiting bottleneck mediated by the antiviral response early in infection that is overcome by the capsid protein ORF2, but not ORF3, allowing for the establishment of persistent viral replication.

## Results

### The HEV ORF2 protein antagonizes both antiviral and inflammatory signaling pathways and interacts with TBK1

Several independent studies have reported that HEV ORF2 and ORF3 antagonize different steps of the cell-intrinsic antiviral response, however, with a variety of different conclusions^22–27^. Therefore, we aimed to mechanistically dissect at which steps ORF2 and ORF3 interfere with antiviral and inflammatory signaling. Importantly, we clearly separated our analysis between IFN induction and IFN signaling as well as systematically compared the roles of ORF2 and ORF3 side by side.

We first focused on sensing and signaling by PRRs. For this mechanistic investigation, we made use of readily available A549-derived cell lines, which harbor a double knockout of RIG-I and MDA5 and show negligible TLR3 expression^29^. In addition to reconstitution with weak transgene overexpression of either MDA5^30^, RIG-I^30^, or TLR3, we ectopically expressed HEV ORF2, ORF3, or GFP in these cells. In GFP-expressing cells, stimulation of the respective PRR with Mengo-Zn virus, Sendai virus (SeV), and poly(I:C) supernatant feeding resulted in the induction of *IFNB1* expression (Fig. 1A-C). Ectopic expression of HEV ORF3 did not significantly alter *IFNB1* expression downstream of RIG-I and MDA5 (Fig. 1A-B) but resulted in a minor enhancement downstream of TLR3 compared to the GFP control (Fig. 1C). In cells expressing ectopic ORF2, *IFNB1* induction was significantly lower downstream of all PRRs compared to GFP-expressing cells (Fig. 1A-C). Not only ORF2 from a chronic HEV genotype (HEV-3) but also from an acute genotype (HEV-1) dampened this response (Fig. 1A-C). By ELISA, we confirmed that ectopic ORF2 expression also decreased IFNβ protein secretion (Suppl. fig. 1A-C). While the presence of ectopic ORF2 dampened the strength of *IFNB1* induction, it did not alter the dynamics of RLR signaling, which we tested by time-resolved RT-qPCR analysis of MDA5-expressing cells electroporated with poly(I:C) (Suppl. fig. 1D-E).

**Figure 1:**
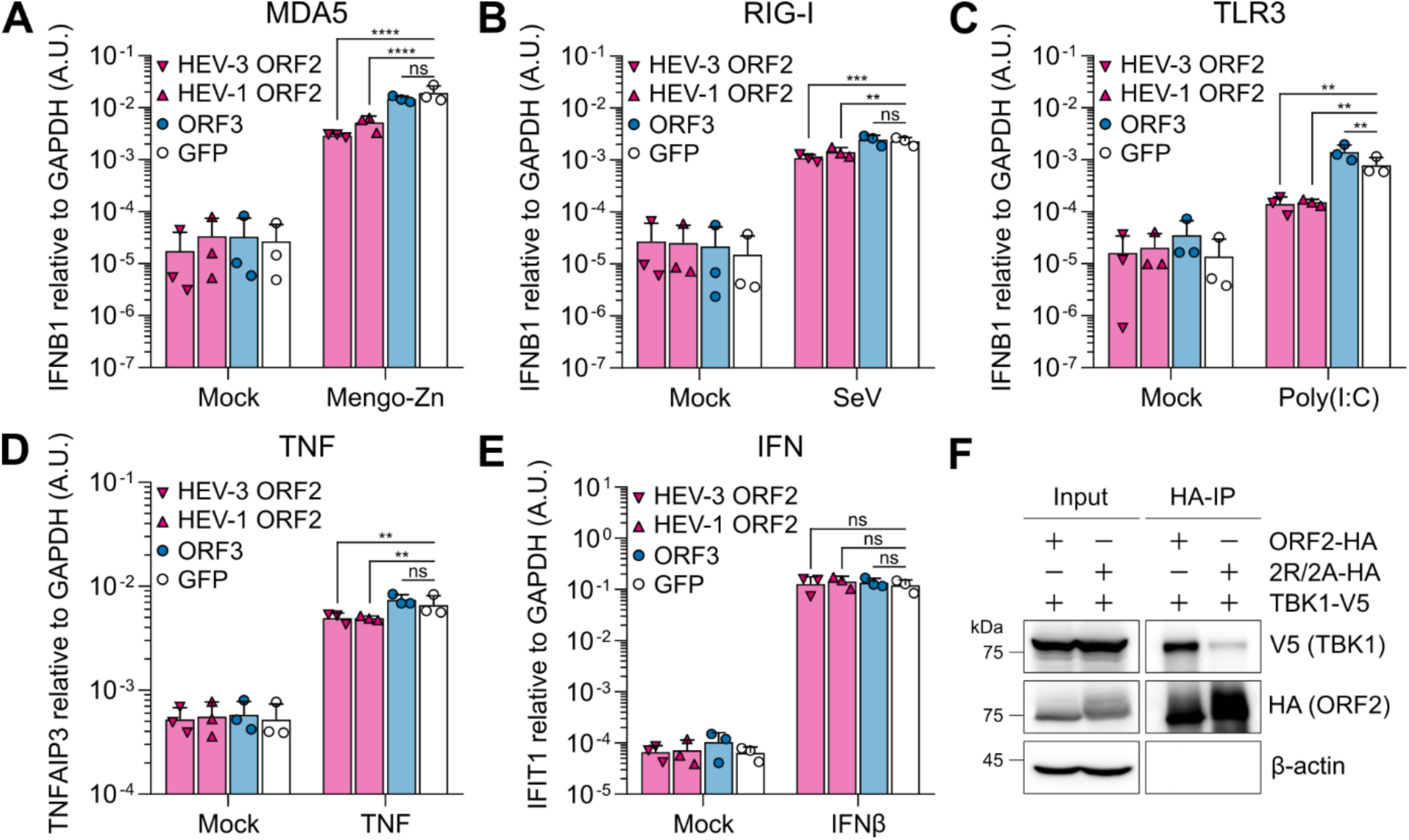
The HEV ORF2 protein antagonizes both antiviral and inflammatory signaling pathways and interacts with TBK1. (A) A549 cells harboring knockouts of the PRRs RIG-I and MDA5 and ectopically expressing a single PRR (MDA5, RIG-I, or TLR3) together with either HEV-3 ORF2, HEV-1 ORF2, ORF3, or GFP were challenged with either Mengo-Zn virus at MOI 1 for 24 h, (B) Sendai virus (SeV) at MOI 0.75 for 4 h, or (C) 50 µg/mL poly(I:C) supernatant feeding for 24 h, as indicated. (D) A549-derived cells ectopically expressing MDA5 were stimulated with 10 ng/mL TNF or (E) 200 IU/mL IFNβ for 8 h. At respective time points, (A-C) *IFNB1*, (D) *TNFAIP3*, and (E) *IFIT1* expression were analyzed by RT-qPCR, each relative to the housekeeping gene *GAPDH* using the 2^-ΔCt^ method. Data show mean ± SD of n = 3 independent biological experiments. Statistical analysis was performed using two-way ANOVA. **: p < 0.01; ***: p < 0.001; ****: p < 0.0001; ns, non-significant. A.U., arbitrary units. (F) HEK293T cells were co-transfected with ORF2-HA or ORF2-2R/2A-HA and TBK1-V5 and lysed 24 h post-transfection. Anti-HA co-IP and WB analysis for TBK1 (anti-V5 staining), ORF2 (anti-HA staining), and β-actin were performed. Shown is a representative blot of n = 3 independent biological experiments.

We further investigated the impact of ORF2 expression on inflammatory and cytokine-dependent antiviral signaling. Upon TNF stimulation, the presence of ectopic ORF2, but not ORF3, reduced the induction of the inflammatory cytokine *TNFAIP3* significantly (Fig. 1D), thus interfering with NF-κB-dependent signaling. On the other hand, we did not observe significant differences in expression of the ISG *IFIT1* upon stimulation with IFNβ in the presence of either ectopic ORF2 or ORF3, suggesting that neither interfered with JAK/STAT signaling (Fig. 1E).

Our results demonstrated that ORF2 interferes with the sensing pathway upstream of IFN induction and with NF-κB-dependent inflammatory cytokine induction downstream of the TNF receptor, similar to previous literature^22–24^. ORF3 did not affect antiviral and inflammatory responses as strongly as proposed previously^25–28^. Importantly, IFN-mediated signaling was not affected by either of the viral proteins. As NF-κB-dependent cytokine induction is particularly important for the crosstalk with immune cells *in vivo*, we continued focusing on the effects of ORF2 on the antiviral IFN/ISG response.

It was previously suggested that ORF2 interacts with the critical adaptor molecule TBK1, likely through the ARM^23^, but a direct interaction through this motif was not analyzed. Here, we confirmed the interaction of HEV-3 ORF2 and TBK1 by co-immunoprecipitation (co-IP) (Fig. 1F). We further found that mutation of the last two arginine residues of the ARM to alanine (2R/2A) decreased the interaction of TBK1 with ORF2 (Fig. 1F). However, this particular mutation abrogates nuclear translocation of ORF2i and increases ORF2 secretion and glycosylation along the secretory pathway^4^. Hence, the pool of cytosolic ORF2i is reduced^4^ and thus, also the likelihood of interaction with TBK1. In agreement with this, the ORF2-2R/2A protein appeared with a light smear in our Western blot (WB) analysis as compared to WT ORF2, suggesting the presence of post-translational modifications such as glycosylation (Fig. 1F).

In an effort to determine other potential interaction motifs between ORF2 and TBK1, we made use of AlphaFold. Repeated modeling identified the last three residues of the LGSA<u>WRD</u> motif (spanning amino acids 83 to 89) at the N-terminus of ORF2 as the only promising, putative interaction motif (Suppl. fig. 2A, 2C-E). However, mutation of these residues to triple alanine did not result in impaired interaction between ORF2 and TBK1, suggesting that this predicted motif is not essential (Suppl. fig. 2B).

Altogether, we systematically validated in a side-by-side comparison that ORF2 but not ORF3 counteracts IFN induction downstream of all relevant PRRs and additionally interferes with NF-κB-mediated signaling. The antagonism of the antiviral response by ORF2 is mediated at least in part by interaction with TBK1, although the precise interaction motif remains elusive.

### The capsid protein ORF2, but not the ORF3 protein, is pivotal for efficient HEV replication in immunocompetent cells

HEV replication in hepatocytes persists in the presence of a sustained cell-intrinsic antiviral response^11,12^. To study how far HEV replication is dependent on ORF2-mediated escape of the antiviral response, we generated a ΔORF2 and a control ΔORF3 mutant of the HEV-3 Kernow-C1/p6 (wild type, WT) strain (Fig. 2A). To obtain ΔORF2, we mutated the two start codons of *ORF2* to abrogate protein expression of all ORF2 isoforms without affecting correct translation of the overlapping *ORF3*, as suggested by Yin *et al.*^5^. Upon electroporation (EPO) into immunocompetent HepG2/C3A cells^11,31^, we observed significantly reduced replication of the ΔORF2 mutant compared to both WT and the ΔORF3 mutant on day 5 and day 7 post-EPO (Fig. 2B). Of note, at least part of the basal HEV RNA detected up to day 3 post-EPO resulted from high levels of incoming, electroporated RNA, as revealed by comparison with a replication-incompetent GNN mutant of the RdRp (Fig. 2B). *IFNL1* (Fig. 2C) and *ISG15* induction (Fig. 2D) upon HEV WT, ΔORF3, but also ΔORF2 replication was similar on day 1 and day 3 post-EPO. However, we observed significantly stronger expression of both antiviral response genes for ΔORF2 on day 5 post-EPO (Fig. 2C-D), coinciding with the decrease in ΔORF2 viral RNA (Fig. 2B). While induction of *IFNL1* and *ISG15* was not significantly different from WT and ΔORF3 on day 7 post-EPO, the 8.6-fold lower levels of ΔORF2 RNA compared to WT RNA (Fig. 2B) indicated a relatively stronger antiviral response induction for ΔORF2 (Fig. 2C-D and Suppl. fig. 3A-B).

**Figure 2:**
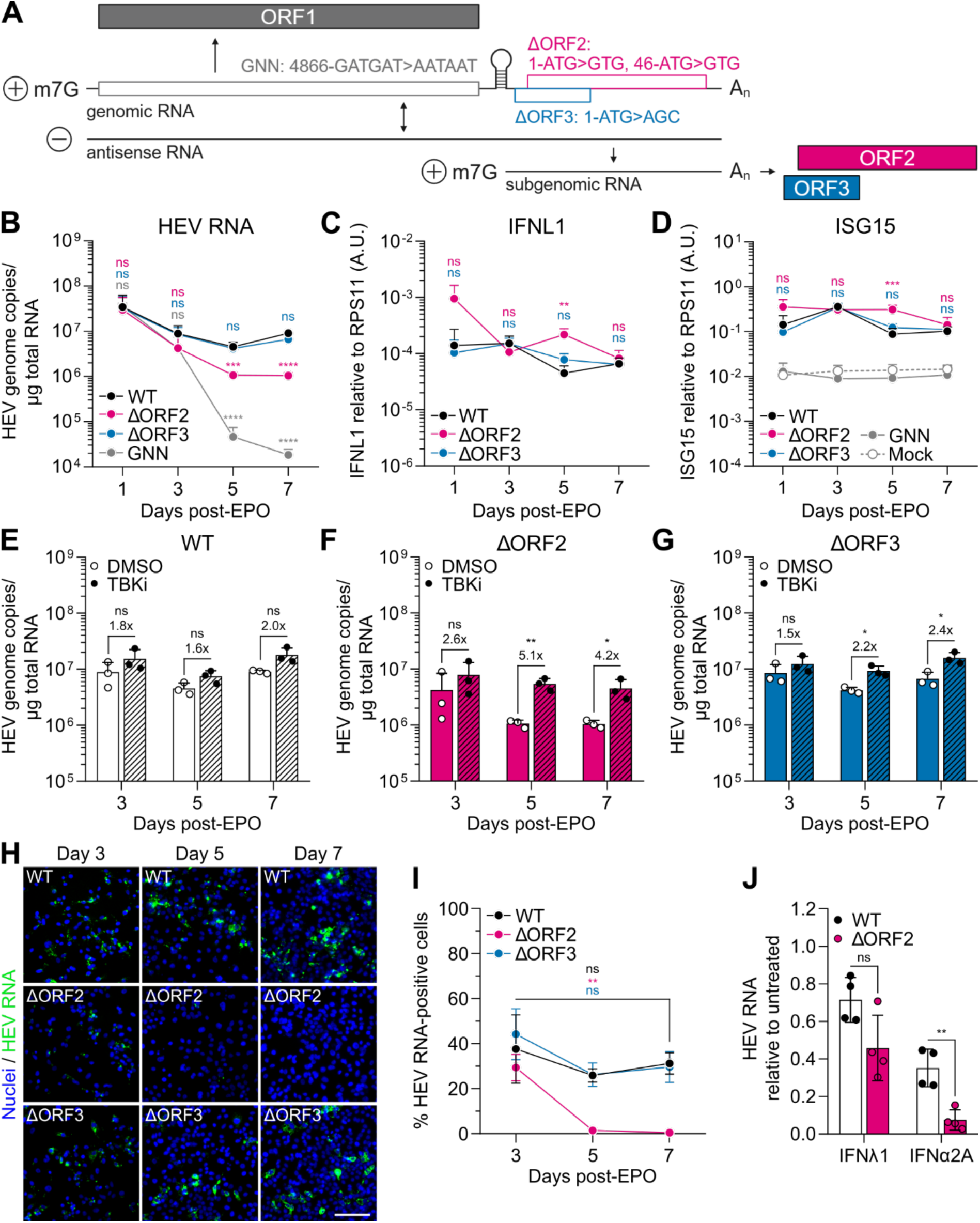
The capsid protein ORF2, but not the ORF3 protein, is pivotal for efficient HEV replication in immunocompetent cells. (A) Schematic representation of the HEV genome and its replication intermediates. Nucleic acid positions of GNN mutation and start codon mutations of ΔORF2 and ΔORF3 mutants are indicated. m7G, 7-methylguanosin cap; A_n_, poly(A)-tail. Created in BioRender. Dao Thi, V. (2025) https://BioRender.com/j85i934 (B) HepG2/C3A cells were electroporated with *in vitro* transcribed (IVT) HEV WT, ΔORF2, ΔORF3, or replication-incompetent GNN RNA and analyzed by RT-qPCR at indicated time points post-EPO for HEV RNA, (C) *IFNL1* expression over the housekeeping gene *RPS11* using the 2^-ΔCt^ method, and (D) *ISG15* expression over the housekeeping gene *RPS11* using the 2^-ΔCt^ method. *IFNL1* expression was undetectable in mock and GNN samples. Data show mean ± SD of n = 3 independent biological experiments. Statistical analysis over WT was performed using one-way ANOVA of each time point independently and is indicated above the respective time points in the corresponding color. **: p < 0.01; ***: p < 0.001; ****: p < 0.0001; ns, non-significant. A.U., arbitrary units. (E-G) Electroporated HepG2/C3A cells from (B) were additionally treated with 6 µM of the TBK1 inhibitor BX795 (TBKi) or respective DMSO vehicle control 48 h prior to the indicated time point post-EPO. HEV RNA was quantified by RT-qPCR and fold changes over DMSO treatment are shown above the respective graphs. Data show mean ± SD of n = 3 independent biological experiments. Statistical analysis was performed using unpaired two-tailed Student’s t-test of each time point independently. *: p < 0.05; **: p < 0.01; ns, non-significant. (H) Electroporated HepG2/C3A cells from (B) were stained for HEV RNA using RNA-FISH at indicated days post-EPO. Images show exemplary images of n = 3 independent biological experiments. Scale bar, 100 µm. (I) The percentages of electroporated HepG2/C3A cells positive for HEV RNA, stained by RNA-FISH in (H), were quantified with ilastik and CellProfiler. Data show mean ± SEM of n = 3 independent biological experiments and 5 images quantified per experiment. Statistical analysis was performed using unpaired two-tailed Student’s t-test of each condition independently and is indicated in the corresponding color. **: p < 0.01; ns, non-significant. (J) Huh7.5 cells were electroporated with IVT HEV WT or ΔORF2 RNA and treated with 10 ng/mL IFNλ1 or 10,000 IU/mL IFNα2A from day 4 to day 7 post-EPO. IFNs were replenished every 24 h. HEV RNA was quantified by RT-qPCR on day 7 post-EPO and normalized over the respective untreated condition. Data show mean ± SD of n = 4 biological repeats from two independent biological experiments. Statistical analysis was performed using unpaired two-tailed Student’s t-test of IFNλ1 and IFNα2A independently. **: p < 0.01; ns, non-significant.

Importantly, we did not observe induction of *IFNL1* (not detectable) or *ISG15* expression (Fig. 2D) upon EPO with the replication-incompetent GNN mutant. We therefore concluded that the incoming RNA and consequently, the full-length single-stranded HEV genome was not sensed by PRRs. Hence, we excluded the possibility that the differences in the antiviral response we observed for ΔORF2 were the results of increased sensing of unpackaged HEV genome due to the missing capsid protein ORF2. Furthermore, we ruled out effects of ORF2 and ORF3 deletion on secondary virus spread because ΔORF3 replicated at WT levels and ORF3 is the protein responsible for secretion of progeny. In addition, newly produced virus particles released into the supernatant are quasi-enveloped and therefore, not highly infectious^32^.

We hypothesized that the antiviral response induced in the absence of ORF2 was strong enough to cause, at least in part, the observed decline in viral replication. In order to address this, we exogenously inhibited IFN and ISG induction by treating the cells with the drug BX795, which inhibits the target of the ORF2 antagonism, TBK1. Prolonged incubation with BX795 can affect cell viability, as TBK1 is also involved in many other cellular processes. To minimize such effects (Suppl. fig. 3C-E), we incubated the cells with BX795 for only 48 h before harvesting. Application of BX795 reduced *IFNL1* and *ISG15* induction in response to viral replication (Suppl. fig. 3F-H). HEV WT replication was not significantly enhanced (Fig. 2E), while ΔORF3 replication showed only a minor increase (Fig. 2G). In contrast, the BX795 treatment resulted in a significant and stronger increase in ΔORF2 replication, especially on day 5 and day 7 post-EPO (Fig. 2F). This implied that the antiviral response had a greater effect on viral replication in the absence of ORF2. To study this observation at the single-cell level, we detected HEV genomes by RNA-fluorescence *in situ* hybridization (RNA-FISH) (Fig. 2H), allowing us to quantify the percentage of HEV-infected cells over time, independent of viral antigen expression. As early as day 3 post-EPO, ΔORF2 RNA signals were weaker compared to WT and ΔORF3 and continued to decline thereafter (Fig. 2H-I). Analogous to our results in Fig. 2B, the decrease in WT- and ΔORF3- positive cells from day 3 to day 5 post-EPO was likely due to the detection of residual incoming RNA on day 3, introduced by the EPO process (Fig. 2I). Overall, the percentages of WT- and ΔORF3-positive cells remained comparable between day 5 and day 7 post-EPO (Fig. 2I). In contrast, we barely detected ΔORF2 RNA at these days as evidenced by a significant decrease compared to day 3. Interestingly, ΔORF2 RNA was still detectable in our bulk RT-qPCR analysis at these time points (Fig. 2B). Importantly, the RNA-FISH assay is based on the hybridization of 20 probe pairs to an *ORF2* target region spanning approximately 1,000 nucleotides^33^, whereas RT-qPCR only amplifies a short sequence stretch of 70 nucleotides. Therefore, it is possible that we detected many degraded ΔORF2 genomes by RT-qPCR at day 5 and day 7 post-EPO.

Next, we aimed to determine whether the absence of ORF2 resulted in increased sensitivity of viral replication to the action of antiviral effectors. To focus solely on the effects of ISGs, we used Huh7.5 cells, which are only weakly responsive to PRR stimulation due to a non-functional RIG-I and negligible basal expression of MDA5^34^, TLR3^34^, and LGP2^35^. Therefore, HEV replication should not induce an antiviral response. Indeed, EPO of Huh7.5 cells with WT and ΔORF2 RNA resulted in comparable viral replication (Suppl. fig. 4A), likely due to the lack of an antiviral response in these cells (Suppl. fig. 4B). Next, we treated WT- and ΔORF2-electroporated Huh7.5 cells four days post-EPO with either IFNλ1 or IFNα2A for 72 h to induce ISG expression. Comparable ISG induction between WT and ΔORF2 further confirmed that ORF2 did not interfere with JAK/STAT signaling (Suppl. fig. 4C-D). Importantly, we detected neither *IFNB1* nor *IFNL1* expression upon IFN treatment of WT- and ΔORF2-electroporated cells. Thus, we could exclude the possibility that the IFN treatment led to an upregulation of PRR expression and consequently, an increased ISG induction in a positive feedback loop due to enhanced sensing of HEV RNA. Nonetheless, we observed that ΔORF2 replication was more strongly inhibited by IFN treatment and the ensuing ISG response than WT replication (Fig. 2J). We therefore concluded that ΔORF2 replication was more sensitive to the antiviral effects of ISGs induced by exogenous IFN treatment, even in the absence of a directly virus-induced antiviral response. We therefore concluded that this additional protective role of ORF2 further contributed to the decrease in viral replication in the absence of ORF2 (Fig. 2B).

Altogether, we found that the presence of ORF2 is critical for efficient HEV replication due to a dampened cell-intrinsic antiviral response. Further, the absence of ORF2 resulted in increased sensitivity of HEV replication to the action of ISGs, suggesting additional protective functions of the ORF2 protein, independently of its direct interaction with TBK1.

### A balance between HEV replication and the antiviral response at an early infection bottleneck is essential for sustained viral replication

We next sought to identify the time point at which the presence of ORF2 is decisive for the fate of HEV replication in authentic infection. To this end, we established a trans-complementation system in hepatoma S10-3 cells ectopically expressing ORF2 to produce ΔORF2 virus particles (Fig 3A), similar to trans-complementation of a subgenomic replicon described previously^36^. Ectopic ORF2 expression was comparable to the protein level reached by viral replication after EPO of WT HEV RNA into naïve S10-3 cells (Fig. 3B). We further confirmed by WB that ORF3 but not ORF2 was expressed upon EPO of ΔORF2 RNA into naïve cells (Fig. 3B). We then harvested infectious ΔORF2 particles from ORF2-expressing S10-3 cell lysates to perform HEV infections.

**Figure 3:**
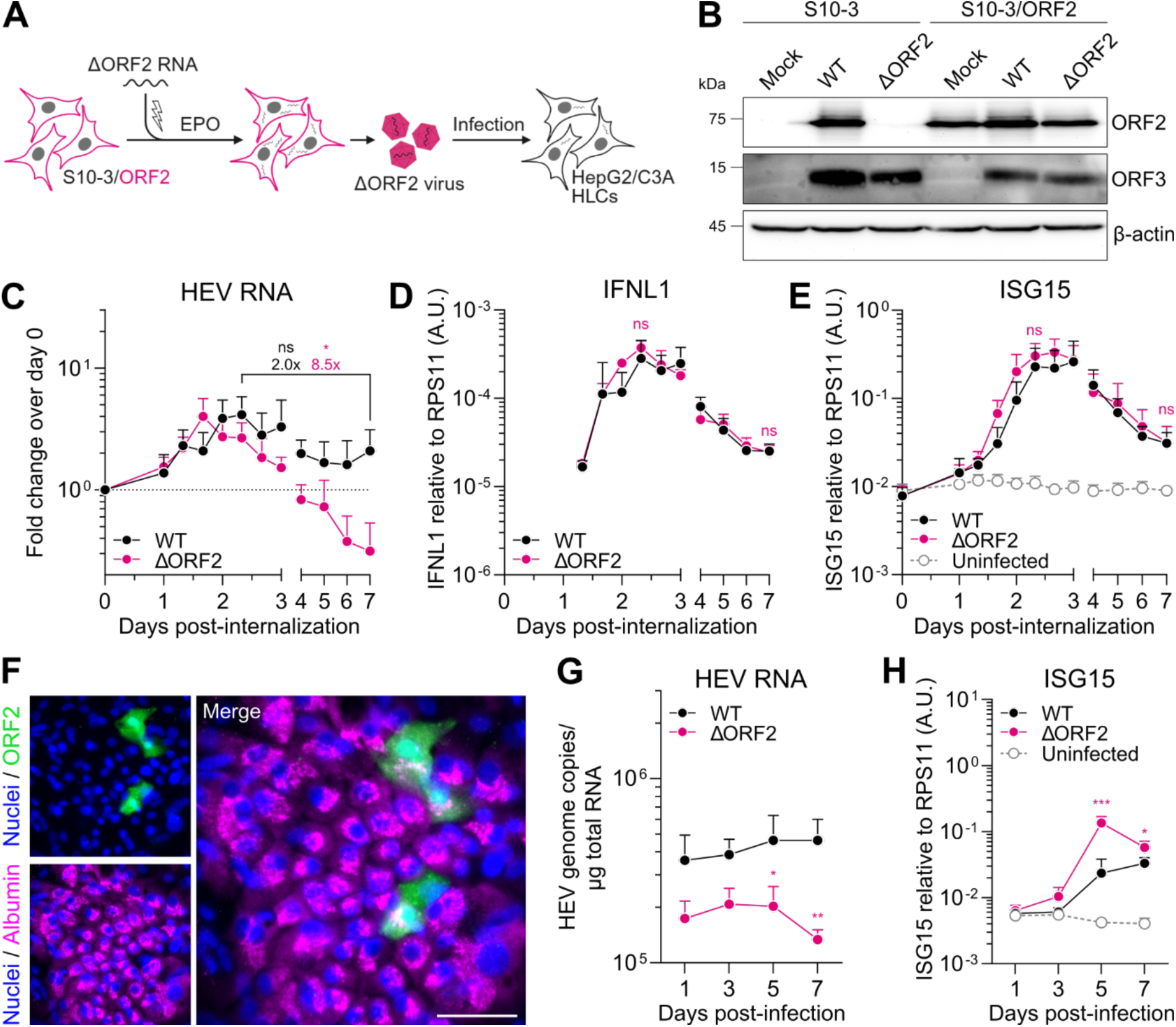
A balance between HEV replication and the antiviral response at an early infection bottleneck is essential for sustained viral replication. (A) Schematic of ΔORF2 trans-complementation. S10-3 cells stably expressing HEV ORF2 were electroporated with ΔORF2 HEV RNA to produce virus particles containing the ΔORF2 genome packaged in capsid ORF2 protein expressed by the producer cell. Trans-complemented ΔORF2 virus was then used to infect HepG2/C3A cells or stem cell-derived HLCs. Created in BioRender. Dao Thi, V. (2025) https://BioRender.com/u54x394 (B) WB analysis of mock, WT-, or ΔORF2-electroporated S10-3 or S10- 3/ORF2 cells for ORF2, ORF3, and β-actin protein expression. (C) Equal genome copies of HEV WT and ΔORF2 virus particles (30 GE/cell) were bound on HepG2/C3A cells for 2 h at 4 °C prior to internalization at 37 °C for 8 h, followed by removal of inoculum (= day 0). RT-qPCR was performed at indicated time points post-internalization to determine fold viral replication over day 0, (D) *IFNL1* expression over the housekeeping gene *RPS11* using the 2^-ΔCt^ method, and (E) *ISG15* expression over the housekeeping gene *RPS11* using the 2^-ΔCt^ method. *IFNL1* expression was undetectable in the uninfected sample. Statistical analysis of fold changes over time (C) and ΔORF2 compared to WT (D-E) are indicated above the respective time points in the corresponding color. Data show mean ± SD of n = 3 independent biological experiments. Statistical analysis was performed using unpaired two-tailed Student’s t-test of the respective conditions or days independently. *: p < 0.05; ns, non-significant. A.U., arbitrary units; n/d, not detectable. (F) Exemplary image of stem cell-derived hepatocyte-like cells (HLCs) infected with HEV WT, fixed, and stained for albumin and ORF2 by immunofluorescence. Scale bar, 50 µm. (G) Stem cell-derived HLCs were infected with equal genome copies of HEV WT and ΔORF2 virus particles (30 GE/cell) and analyzed by RT-qPCR for HEV RNA and (H) *ISG15* expression over the housekeeping gene *RPS11* using the 2^-ΔCt^ method. Data show mean ± SD of n = 4 biological replicates of two independent HLC differentiations. Statistical analysis of ΔORF2 compared to WT was performed using unpaired two-tailed Student’s t-test of each time point independently. *: p < 0.05; **: p < 0.01; ***: p < 0.001.

Next, we aimed to study the replication dynamics after cell entry in HepG2/C3A cells in a time-resolved manner. We synchronized the infection at equal MOI (30 genome equivalents (GE)/cell) of both WT and ORF2-trans-complemented HEV particles by a binding step for 2 hours (h) at 4 °C and internalization for 8 h at 37 °C after which the inoculum was removed, referred to as day 0 post-internalization. We analyzed viral replication and the antiviral response every 8 h between day 1 and day 3 post-internalization and every day (i.e., every 24 h) thereafter by RT-qPCR. Despite infection with the same number of GEs, ΔORF2 HEV RNA levels detected after internalization were always approximately 2.4-fold lower than WT HEV RNA (Suppl. fig. 5A). This indicated a lower specific infectivity of the trans-complemented ΔORF2 virus particles, potentially due to less efficient progeny assembly. For easier comparison, we therefore normalized HEV RNA over day 0 post-internalization.

Replication of WT and the ΔORF2 mutant peaked at 48±8 h post-internalization, followed by a decline (Fig. 3C). Interestingly, while the HEV WT replication leveled off between 56 h and day 7 post-internalization (2.0-fold decrease), ΔORF2 replication continued to decrease significantly by 8.5-fold between those two time points. Following the onset of viral replication, we observed an increase in *IFNL1* (Fig. 3D) and *ISG15* expression (Fig. 3E), which peaked at 56 h and 64 h, respectively. Following the decrease in viral replication, *IFNL1* and *ISG15* levels also declined. *IFNL1* and *ISG15* expression for ΔORF2 was not significantly different from WT at 56 h and on day 7 post-internalization (Fig. 2D-E). However, considering the fact that ΔORF2 RNA significantly decreased until day 7 compared to WT RNA (Fig. 2C), the comparable *IFNL1* and *ISG15* induction between WT and ΔORF2 suggested a relatively stronger antiviral response in ΔORF2 infection (Suppl. fig. 5B-C). Importantly, we did not detect an antiviral response following HEV internalization (day 0) (Fig. 3D-E). This is in agreement with our earlier finding that the incoming RNA and thus, the full-length single-stranded HEV RNA is not being sensed by the cells. Next, we aimed to validate our findings in a more physiologically relevant hepatocellular system. To this end, we used human pluripotent stem cell-derived hepatocyte-like cells (HLCs) which are permissive to HEV infection^12,32^ (Fig. 3F). We infected HLCs overnight at equal MOI (30 GE/cell) with HEV WT and ΔORF2 virus particles produced by trans-complementation. Similar to our results in infected HepG2/C3A cells, ΔORF2 RNA was lower than WT on day 1 post-infection, likely due to a lower specific infectivity as discussed above. Nonetheless, we observed decreased ΔORF2 replication over time (Fig. 3G), accompanied by a significantly stronger induction of *ISG15* compared to HEV WT on day 5 and day 7 post-infection (Fig. 3H).

Overall, our observations suggest that the presence of the ORF2 protein is critical for establishing an equilibrium between viral replication and the antiviral response early in infection, allowing HEV to replicate in different immunocompetent hepatocellular systems.

### scRNA-seq analysis reveals globally dampened and partly divergent ISG responses in HEV-infected cells and bystanders in the presence of ORF2

Expression of ISGs can be induced immediately downstream of viral recognition by PRRs (reviewed in^9^). In addition, secreted IFN can lead to amplified ISG induction in an auto- and paracrine manner. In HEV infection, it remains unclear whether IFN and ISG responses observed in bulk analyses such as RT-qPCR are derived from infected cells, uninfected bystanders, or possibly both. ISG responses from uninfected bystanders only could be induced by low levels of IFN secretion from infected cells. Alternatively, viral RNA transfer to neighboring cells via extracellular vesicles could lead to IFN and ISG induction in bystanders, similar to what has been described for HCV and other viruses (reviewed in^37^).

To identify the cellular origin of the antiviral response in HEV infection and assess potential differences in the response in the absence of ORF2, we performed single-cell RNA-sequencing (scRNA-seq) using microfluidics-based 3’-targeted 10x Genomics. To this end, we infected HepG2/C3A cells with HEV WT and trans-complemented ΔORF2 virus particles in a synchronized manner as described above and analyzed the cells at 56 h post-internalization. This time point represents the antiviral response-mediated bottleneck we identified to be essential for persistent replication in the presence of ORF2 (Fig. 3C-E). Since the ΔORF2 virus replicates to almost undetectable levels at later time points, we only analyzed HEV WT-infected cells on day 7 post-infection.

We first performed a gene set enrichment analysis of the HEV WT- and ΔORF2-infected samples compared to the uninfected sample at 56 h, using the hallmark gene sets of the Human MSigDB Collections^38^. Interferon alpha and gamma responses were among the most significantly upregulated gene sets, in both HEV WT and ΔORF2 infection, highlighting that the global response was dominated by differential expression of ISGs (Fig. 4A-B). Enrichment of oxidative phosphorylation and glycolysis gene sets indicated increased metabolic activity of the infected samples. Analysis of HEV WT infection on day 7 showed that the enriched gene sets were comparable over time and interferon-related responses were still prominently enriched (Fig. 4C).

**Figure 4:**
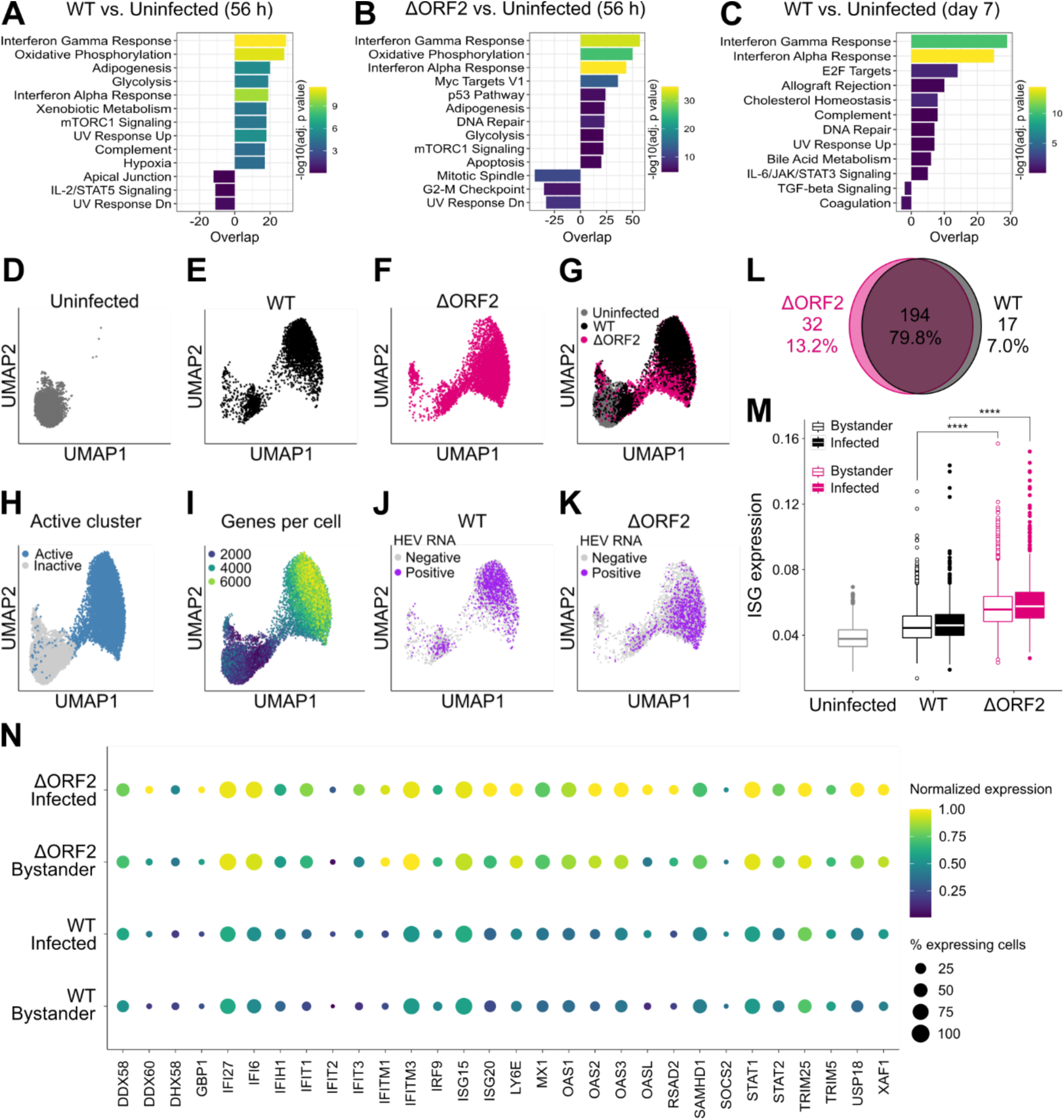
scRNA-seq analysis reveals globally dampened and partly divergent ISG responses in HEV-infected cells and bystanders in the presence of ORF2. (A) Uninfected, WT-infected and ΔORF2-infected HepG2/C3A cells were harvested and processed by microfluidics-based 3’-targeted 10x Genomics at 56 h post-synchronized infection. Uninfected and WT-infected HepG2/C3A cells were also analyzed on day 7 post-synchronized infection. Gene set enrichment analysis comparing WT-infected with uninfected sample at 56 h post-infection, (B) ΔORF2-infected with uninfected sample at 56 h, and (C) WT-infected with uninfected sample on day 7 post-infection based on the hallmark gene sets of the hallmark gene sets of the Human MSigDB Collections^38^ was performed. (D) UMAP projections of uninfected, (E) WT-infected, or (F) ΔORF2-infected HepG2/C3A cells harvested at 56 h post-synchronized infection. UMAPs were generated by clustering of all cells based on a list of approximately 400 ISGs published previously by Schoggins *et al.*^39^ (G) Combined UMAP projections of uninfected, WT-, and ΔORF2-infected samples, colored by condition. (H) Cells with upregulated ISG expression are highlighted in UMAP projections of uninfected, WT-, and ΔORF2-infected samples in blue, representing the active cluster. (I) UMAP projection of uninfected, WT-, and ΔORF2-infected samples, highlighting the number of genes detected per cell in all samples. (J) UMAP projections of the WT-infected and (K) ΔORF2-infected samples with indicated binarized HEV RNA counts in purple. (L) Differentially expressed ISGs in the active clusters of WT- and ΔORF2-infected samples were compared and plotted in a Venn diagram. Absolute numbers and percentages of shared and distinct differentially expressed ISGs are shown. (M) Global changes in the ISG response were analyzed by comparison of normalized ISG expression across all samples (Uninfected, WT, and ΔORF2), based on a list of approximately 400 ISGs. WT- and ΔORF2-infected samples were split into actively infected cells, defined by detection of at least one HEV RNA copy per cell, and uninfected bystanders. Statistical analysis was performed using a two-sided Wilcoxon rank sum test. ****: p < 0.0001. (N) Out of the overlapping, differentially expressed ISGs between the active clusters of WT- and ΔORF2-infected samples, 30 ISGs were selected. Normalized expression and the percentage of cells were plotted for WT- and ΔORF2-infected samples, split by actively infected cells (at least one HEV RNA count per cell) and uninfected bystanders.

Next, we clustered the uninfected, HEV WT-, and ΔORF2-infected samples at 56 h based on a list of approximately 400 ISGs published previously by Schoggins *et al.*^39^ (Fig. 4D-G). This revealed two major subpopulations in the infected samples (Fig. 4E-F), also on day 7 (Suppl. fig. 6A-C). We concluded that the cells located in the cluster overlapping with the uninfected sample were non-responding cells, characterized by unchanged ISG expression. The cells located in the second cluster showed differential ISG expression compared to the non-responding cells and were thus classified as responders, from here on called the active cluster (Fig. 4H). In agreement with the globally enhanced metabolic activity (Fig. 4A-B), this cluster appeared to be more transcriptionally active, as indicated by an increased number of expressed genes (Fig. 4I) as well as an overall increased number of transcripts (Suppl. fig. 6E). Within the WT- and ΔORF2-infected samples, 28.1% and 17.2% of cells were identified as HEV RNA-positive, respectively (Suppl. fig. 6F). Interestingly, we observed that most infected cells of both WT- (Fig. 4J) and ΔORF2-infected samples (Fig. 4K) were located in the active cluster, together with uninfected bystanders. We therefore concluded that both actively infected cells and uninfected bystanders responded by upregulated ISG expression. Although detectable by qRT-PCR (Fig. 3D), we could not detect type III IFN gene expression in the infected samples via scRNA-seq, likely due to low numbers of IFN transcripts per cell. Therefore, we could not identify whether only infected cells or also uninfected bystanders were the source of IFN. Nonetheless, we could demonstrate that the ISG response was not fully suppressed in actively infected cells by the ORF2 antagonism.

Compared with the infected cells in the active cluster, the few infected cells located within the inactive cluster had lower HEV RNA counts (Suppl. fig. 6G). These cells might be at an earlier stage of infection due to a delayed onset of replication or due to secondary infection, which, however, should be limited at this early time point. Finally, not only uninfected bystanders but also HEV-infected cells still showed enhanced ISG expression and localized to the active cluster on day 7 post-HEV WT infection (Suppl. fig. 6D). This provided further evidence that the establishment of an equilibrium between HEV replication and the antiviral response early in infection is essential for persistent viral replication in an antiviral environment.

Next, we sought to examine the specific ISG signatures (i.e., the set of ISGs) induced upon WT and ΔORF2 infection. We compared the respective lists of differentially expressed ISGs in the active clusters of WT and ΔORF2 infection and found that out of 243 differentially expressed ISGs, 79.8% overlapped (Fig. 4J and Suppl. table 2). This was further substantiated by identification of ISGs specifically upregulated in actively infected cells, defined by detection of at least one HEV RNA copy per cell, or bystanders, defined by the absence of HEV RNA, in the active clusters of WT- (Suppl. fig. 7A) and ΔORF2-infected samples (Suppl. fig. 7B). Actively ΔORF2-infected cells expressed a larger subset of differing ISGs compared to WT-infected cells (Suppl. fig. 7A). We therefore concluded that the ISG signatures induced by WT and ΔORF2 infection are not entirely identical.

Next, we compared the strength of ISG induction in actively infected cells and uninfected bystanders within the active cluster. We observed a globally stronger ISG response in ΔORF2- infected cells and bystanders compared to WT infection (Fig 4I), indicating overall increased IFN production and secretion, despite remaining undetectable by scRNA-seq. Consulting the list of overlapping, significantly upregulated ISGs identified in Fig. 4L (and Suppl. table 2) and previous literature on the upregulation of ISGs in HEV infection of different hepatocellular systems and in chimpanzees^10–12,40^, we selected a subset of ISGs for further comparison of infected cells and bystanders. These common differentially expressed genes further substantiated a globally stronger ISG response in both infected cells and uninfected bystanders in the active cluster of the ΔORF2-infected sample compared with the respective cell populations of the WT-infected sample (Fig. 4N).

Overall, we identified both actively infected cells and uninfected bystanders to be the cellular origins of ISG responses at early and later time points of HEV infection. Early in infection, the absence of ORF2 results in induction of a globally stronger and partially divergent ISG signature in infected cells. In addition, increased autocrine and paracrine IFN signaling, when not dampened by the presence of ORF2, leads to enhanced ISG responses in both infected cells and bystanders.

## Discussion

HEV replication has been shown to persist despite a sustained cell-intrinsic antiviral response. However, an in-depth understanding of the underlying mechanisms and the active contribution of a viral antagonism is missing. In this study, we analyzed authentic HEV infection with mutants lacking expression of individual viral proteins at single-cell resolution. We identified a replication-limiting bottleneck imposed by the antiviral response where the presence of ORF2 and thus, likely its interaction with the adaptor molecule TBK1 determine whether viral replication declines or equilibrates with the antiviral response (summarized in Fig. 5).

**Figure 5:**
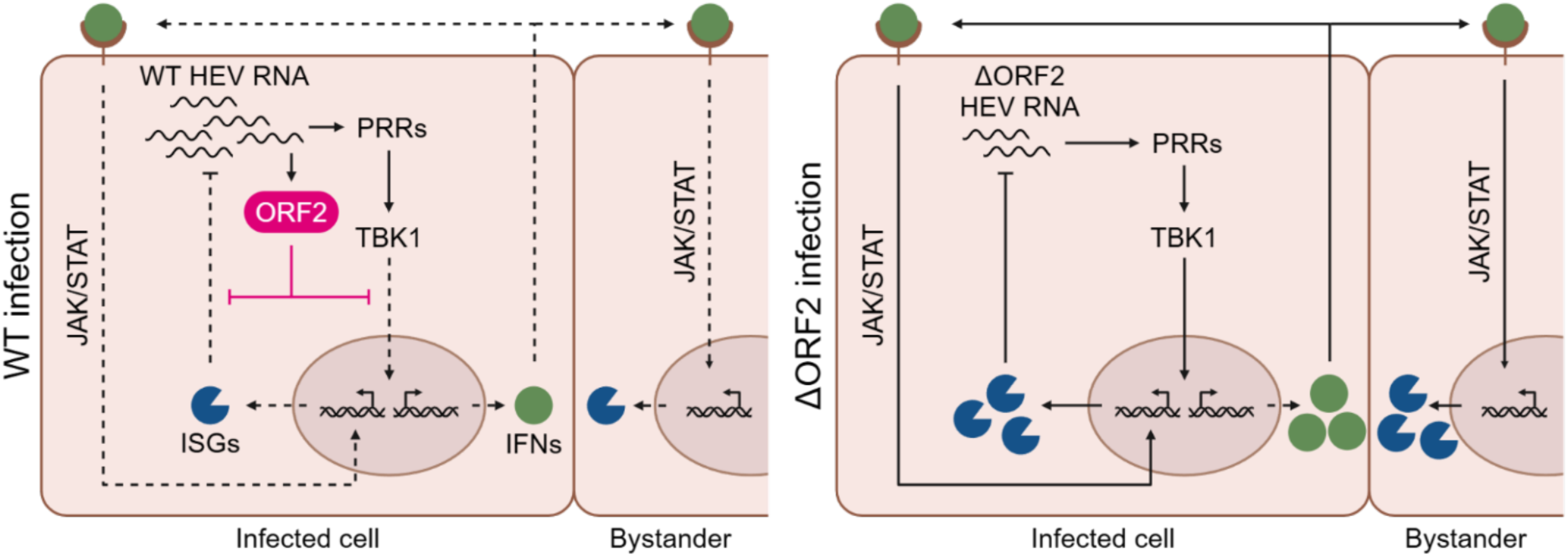
Working model of HEV ORF2 enabling persistent viral replication by dampening the cell-intrinsic antiviral response. ORF2 directly interferes with IFN induction downstream of PRR recognition of HEV RNA in WT-infected cells, at least in part through direct interaction with TBK1. Consequently, ISG induction downstream of PRR signaling as well as IFN secretion and thus, autocrine and paracrine ISG induction downstream of the IFN receptor is dampened in infected cells and bystanders, respectively. Additionally, ORF2 shields HEV RNA from the antiviral functions of ISGs. In ΔORF2-infected cells, the lack of ORF2 results in stronger ISG induction downstream of PRR signaling as well as enhanced IFN secretion, leading to a stronger autocrine and paracrine ISG induction downstream of IFN signaling. Viral replication is consequently dampened as a result of the actions of antiviral effectors. Created in BioRender. Dao Thi, V. (2025) https://BioRender.com/h01q004

### Single-cell analysis uncovers the role of the multifunctional capsid protein for HEV persistence amidst an antiviral environment

In the present study, we have shown that the capsid protein ORF2 is a critical viral determinant of HEV replication efficiency. In absence of the ORF2 protein, a stronger cell-intrinsic antiviral response is induced. Importantly, we ruled out that the full-length, single-stranded HEV genome was sensed because the replication-incompetent GNN mutant did not induce IFN or ISG expression (Fig. 2C-D). Otherwise, sensing of accumulating unpackaged HEV genome due to the missing capsid protein ORF2 could have explained the enhanced antiviral response. A previous study has suggested that the 3’-untranslated region (UTR) of single-stranded HEV RNA is sensed by RIG-I^41^. However, as the authors used transfection of short RNA fragments, which are ideal PAMPs for RIG-I, we hypothesize that this might be rather irrelevant in authentic HEV infection and that double-stranded replication intermediates might be sensed instead.

Investigation of the cell-intrinsic antiviral response at the single-cell level revealed that both infected cells and uninfected bystanders are the source of the ISG response in HEV infection. The ISG response in both cell types was globally stronger in the absence of ORF2 (Fig. 4M-N). Thus, the ORF2-mediated viral antagonism does not completely suppress antiviral responses within infected cells. Instead, it reduces ISG and presumably IFN induction within infected cells and consequently further ISG induction via autocrine and paracrine IFN signaling. Absence of ORF2 further allows expression of a partially divergent ISG subset. It remains to be clarified whether these ISGs might be particularly potent in restricting viral replication. Overall, ORF2 contributes to dampening of the antiviral responses below a critical threshold, allowing viral replication and the antiviral response to reach an equilibrium where they can coexist. We propose that this equilibrium is a prerequisite for persistent HEV infection. Future studies are needed to determine whether IFN is produced only by infected cells or also by uninfected bystanders, and to characterize the role of secreted ORF2 isoforms in influencing paracrine immune signaling.

HEV ORF2 appears to be a multifaceted protein with a wide range of functions affecting different host pathways. In addition to its direct involvement in immune evasion downstream of PRR recognition, we speculate that the capsid protein ORF2 may also exert additional, protective functions. Here, we demonstrated that ORF2 is essential for maintaining the relative resistance of HEV to type I and type III IFN treatment, even in the absence of a virus-induced antiviral response (Fig. 2J). Importantly, we were able to exclude a direct inhibition of IFN signaling by ORF2 (Fig. 1E). Therefore, we concluded that the capsid protein additionally contributes to the shielding of replicating RNA from the action of ISGs and potentially also from recognition of double-stranded replication intermediates by PRRs.

Most positive-sense RNA viruses replicate in the cytosol and restructure their host cell by inducing host membrane-derived replication compartments (reviewed in^42^). While such replication organelles have not yet been reported for HEV, several studies have suggested that different compartments of the endolysosomal system or the ER-Golgi intermediate compartment (ERGIC) could be involved^43–46^. ORF2 was frequently detected at these proposed replication sites. Therefore, it may be involved in either remodeling or redirecting cellular structures to help create the replication site, thereby potentially contributing to protection of viral RNA from the antiviral effects of induced ISGs.

Despite the relatively low abundance of ORF1 and ORF3 proteins compared to ORF2 protein in authentic infection^47^, we cannot exclude the possibility that these proteins have additional potential to interfere with antiviral signaling and contribute to viral persistence. Ectopically expressed ORF3 may interact with components of the antiviral response as reported by others^25–27^. In this study however, we did not observe any striking effects of ORF3 on viral replication and antiviral responses. Furthermore, the processing of the ORF1 polyprotein in authentic infection remains controversial (reviewed in^48^). Therefore, physiologically relevant contributions of the previously reported antagonisms by ORF1 to the overall persistence phenotype remain to be confirmed.

### The direct interaction of HEV ORF2 with TBK1 is critical for antagonizing replication-limiting responses

There is no convincing evidence for a viral protease encoded by the HEV genome. Indeed, unlike HCV^49^ or HAV^50^, HEV does not cleave MAVS^11^, a key adaptor molecule in the induction of cell-intrinsic antiviral responses. Instead, HEV appears to have developed other counteractive strategies. In agreement with previous literature^22,23^, we found that the HEV ORF2 protein interacts directly with TBK1 to dampen IFN induction. This could be directly involved in impairing the establishment of an infection-limiting antiviral state. Here, we identified an early time point at which the presence of ORF2 is critical for the progression of authentic HEV infection, facilitating a balance between viral replication and the antiviral response within immunocompetent cells.

Lin and colleagues^23^ previously suggested that the ARM of ORF2 is responsible for the inhibition of TBK1-induced phosphorylation of IRF3. Indeed, we could show a reduced interaction between ORF2 and TBK1 upon mutation of the ARM. However, this motif has also been identified as the master regulator of the ORF2 isoforms^4^. Therefore, mutating the ARM could also indirectly affect interference with the antiviral response, as enhanced secretion of glycosylated ORF2 along the secretory pathway results in decreased cytosolic and nuclear availability of ORF2 to interact with TBK1 and other unidentified factors. Furthermore, we found that ORF2 also antagonizes NF-κB-dependent cytokine induction (Fig. 1D) and is hence likely to interfere with additional components other than TBK1. This observation is in agreement with the previously published interaction of ORF2 with beta-transducin repeat containing protein (βTRCP)^24^, which resulted in reduced proteasomal degradation of the inhibitory complex of NF-κB. Additionally, nuclear translocation of ORF2 has been proposed to affect inflammatory cytokine induction^4^. Considering the numerous suggested ORF2 antagonisms by us and others, we believe that viral persistence is likely based on combinatorial interference of ORF2 with different pathways.

Due to the apparent multifunctionality of the ORF2 protein, individual functions are difficult to untangle if specialized domains remain unknown. Indeed, our attempts to predict any putative interaction sites between ORF2 and TBK1 with AlphaFold remained unsuccessful. The only promising prediction was localized to the N-terminus of ORF2, however, this portion of the ORF2 protein appears to be intrinsically disordered^51^. While AlphaFold has demonstrated success in identifying binding regions between other viral proteins and host factors^52^, this particular interaction highlights the challenges of accurately predicting interactions involving disordered protein regions^53,54^. This complexity could stem from several factors: AlphaFold heavily relies on co-evolutionary signals, which might be less pronounced in this virus-host interaction. Instead, the ORF2-TBK1 interaction may rather be driven by physico-chemical properties that are not explicitly considered in AlphaFold’s predictions. Finally, the N-terminus of ORF2 may adopt a specific structure upon interaction with TBK1, as is the case for many intrinsically disordered proteins (reviewed in^55^). High-resolution structural studies of TBK1-ORF2 complexes may therefore be required to further identify the interacting regions.

### Innate immunity as a potential determinant of HEV chronicity and species tropism

Infections with different HEV genotypes result in distinct clinical outcomes. While HEV-1 infections are acute and self-limiting, HEV-3 infections can become chronic in immunocompromised individuals. The determinants of these intergenotypic differences remain to be elucidated, but innate and adaptive immune responses are likely critical parameters (reviewed in^16^).

We have previously shown that infection with an HEV-1 isolate resulted in a stronger IFN and ISG response than infection with an HEV-3 isolate. Interestingly, HEV-1 but not HEV-3 replication appeared to decrease thereafter^12^. Why HEV-1 induces a stronger response and whether this contributes to the self-limiting course of infection observed in humans and in HEV- 1-infected chimpanzees^10^ should be investigated in the future.

On the other hand, a moderate but prolonged cell-intrinsic IFN response induced by HEV- 3 could help to induce tolerance and dampen the effect of the stronger type I IFN response elicited by professional immune cells such as plasmacytoid dendritic cells (pDCs). In agreement, we and others have demonstrated a remarkable resistance of HEV replication to exogenous IFN once replication is fully established^11,13–15^. Continuous activation of JAK/STAT signaling upon HEV replication has been shown to render the cells refractory to exogenous IFN stimulation^11^. A recent publication further suggested that ORF2 isoforms modulate detection of HEV-infected cells by pDCs, indicating that this cell type might be contributing to HEV control *in vivo*^31^. Not only do pDCs produce IFN and other cytokines to activate innate immune cells, but they also capture and process antigens to initiate adaptive T cell responses. Consequently, a reduced T cell activation could be beneficial for establishing HEV persistence *in vivo*. Studying the crosstalk with immune cells will be pivotal for identifying determinants of chronicity.

Our results showed no difference in the potential of ORF2 from acute and chronic genotypes to interfere with IRF3- and NF-κB-mediated signaling (Figure 1), which is consistent with previous studies^22,23^. However, this could play a role in the genotype-dependent species tropism of HEV. While HEV-1 is limited to primates and human infection, HEV-3 can infect a wide range of animals and is mainly transmitted zoonotically to humans. Interaction of ORF2 with TBK1 could potentially represent a prime example of the ongoing arms race between virus and host, driving the co-evolution of viral evasion strategies and host defenses. Therefore, future research should investigate the differences in the ability of HEV-1 and HEV-3 ORF2 to counteract TBK1 from different HEV-3 host species, including pig or rabbit. Such studies might contribute to the identification of the determinants of HEV species tropism.

Many open questions remain about the mechanisms of HEV persistence. In this study, we have shown that the HEV capsid protein ORF2 mediates a viral antagonism affecting both antiviral and inflammatory signaling. ORF2 is essential at a replication-limiting bottleneck early in infection mediated by the antiviral response, which is decisive for establishment of persistent viral replication. We have also laid the groundwork for future studies to focus on the role of intergenotypic differences in the antiviral response and crosstalk with immune cells. This could eventually lead to a better understanding of the determinants of acute and chronic manifestations of HEV infection, which will be valuable in identifying novel therapeutic regimens.

## Materials and Methods

### Standard cell culture

The human hepatoma cell lines HepG2/C3A (ATCC HB-8065), Huh7.5 (a kind gift from Charles Rice, The Rockefeller University), S10-3 (a kind gift from Suzanne Emerson, NIH), and derived S10-3/ORF2 as well as the human embryonic kidney cell line HEK293T (ATCC CRL- 3216) were cultured in Dulbecco’s Modified Eagle Medium (DMEM, Gibco, high glucose, GlutaMAX supplement) with 10% fetal bovine serum (FBS, Capricorn) and 1% penicillin/streptomycin (pen/strep), referred to as complete DMEM (cDMEM). HepG2/C3A cells were grown on collagen-coated cell culture vessels. A549-derived cell lines were cultured in DMEM (Gibco, high glucose) with 10% FBS (Pan Biotech), 1% pen/strep, and 1% non-essential amino acids (NEAA). Cell lines with ectopic protein expression were produced by standard lentiviral transduction, selected, and continuously cultured under respective antibiotic selection pressure. Cells were maintained at 37 °C in 95% humidity and 5% CO_2_ atmosphere. All cell lines used in this study routinely tested negative for mycoplasma.

### Stimulation of PRR-overexpressing A549-derived cell lines

1x10^5^ A549-derived cell lines were seeded in 24-well plates. The next day, cells were infected with Mengo-Zn virus^56^ (MOI 1) for 24 h, Sendai virus (SeV, MOI 0.75; prepared from allantoic fluid of embryonated chicken eggs, a kind gift from Rainer Zawatzky, German Cancer Research Center (DKFZ), Heidelberg, Germany) for 4 h, or supernatant-fed with high molecular weight (HMW) poly(I:C) (InvivoGen) at 50 µg/mL for 24 h in DMEM containing 2% FBS, 1% NEAA, and 1% pen/strep. Stimulation with TNF (Abcam) at 10 ng/mL or IFNβ (R&D Systems) at 200 IU/mL was performed in cDMEM plus 1% NEAA for 8 h. At respective time points, cells were lysed for RNA extraction with Monarch Total RNA Miniprep Kit (New England Biolabs).

### Generation of HEV ΔORF2, ΔORF3, and GNN mutants

Mutants were generated in the plasmid pBlueScript SK(+) encoding the HEV-3 Kernow- C1/p6 sequence (pBSK-HEV-p6, GenBank accession number: JQ679013.1) by site-directed mutagenesis. Mutations were introduced by overlap extension PCR with Phusion or Q5 polymerase (New England Biolabs) using the primers listed in Suppl. table 1, followed by restriction enzyme digestion (New England Biolabs), ligation with T4 DNA Ligase (New England Biolabs), transformation of JM109 competent cells (Promega), and validation by Sanger DNA sequencing.

### In vitro transcription and EPO of HEV WT, ΔORF2, ΔORF3, and GNN mutants

Plasmids were linearized with MluI and RNA was *in vitro* transcribed using the mMESSAGE mMACHINE T7 kit (Invitrogen). 4x10^6^ HepG2/C3A cells were electroporated at 270 V and 975 µF with 10 µg of IVT HEV RNA in cytomix (120 mM KCl, 0.15 mM CaCl2, 10 mM KPO_4_, 25 mM HEPES, 2 mM EGTA, and 5 mM MgCl_2_), supplemented with adenosine triphosphate (ATP) and glutathione (GT). After resuspension in cDMEM, HepG2/C3A cells electroporated with HEV WT or mutants were mixed 1:1 with mock-electroporated cells. 2x10^5^ cells/well were seeded on 24-well plates for analysis by RT-qPCR and 1x10^5^ cells/well were seeded on 48-well plates for analysis by RNA-FISH. One day post-EPO, all wells were washed twice with PBS to remove inoculum and cDMEM was replaced. Cell lysates were harvested for RNA extraction using the Universal RNA kit (Roboklon) according to manufacturer’s instructions. On day 1, day 3, and day 5 post-EPO, cDMEM was replaced and 6 µM TBK1 inhibitor BX795 (InvivoGen) or corresponding DMSO vehicle control was added to the wells dedicated to cell lysis 48 h later. Samples for RNA-FISH were fixed at respective time points with 4% paraformaldehyde (PFA).

### EPO of Huh7.5 cells and IFN treatment

Huh7.5 cells were electroporated with HEV WT or ΔORF2 IVT RNA as described above. For time-course analysis, electroporated Huh7.5 cells were resuspended in cDMEM, seeded in 24-well plates, and harvested for RT-qPCR analysis at respective time points. For IFN treatment, cells were plated in a T75 cell culture flask after EPO. On day 3 post-EPO, cells were reseeded to 24-well plates at approximately 80% confluency. One day later, cells were treated with 10,000 IU/mL IFNα2A or 10 ng/mL IFNλ1. IFNs were replenished daily until cells were lysed for RNA extraction with the Universal RNA kit (Roboklon) and RT-qPCR on day 7 post-EPO.

### ΔORF2 virus production by trans-complementation

S10-3/ORF2 cells were electroporated with IVT ΔORF2 RNA and expanded on day 3 post-EPO. Intracellular nHEV particles were harvested 7 days post-EPO from cell lysates by four repeated freeze-thaw cycles and pelleted by ultracentrifugation at 28,000 rpm for 3 h through a 20% sucrose cushion. HEV WT and ΔORF2 genome copies were determined by RT-qPCR following TRIzol extraction from the resuspended virus prep.

### Synchronized, time-resolved HEV infection

6x10^4^ HepG2/C3A cells were seeded on collagen-coated 24-well plates. The next day, cells were infected with equal genome equivalents (GEs) of HEV WT or ΔORF2 virus (30 GE/cell). Virus was bound at 4 °C for 2 h in Minimum Essential Medium (MEM, Gibco), supplemented with 10% FBS and 1% pen/strep, followed by internalization at 37 °C for 8 h. After internalization, inoculum was removed, cells were washed twice with PBS, and cDMEM was added. Samples for RT-qPCR were lysed at respective time points with TRIzol for RNA extraction.

### 3’-targeted 10x Genomics and Illumina sequencing

8x10^4^ HepG2/C3A cells were seeded on collagen-coated 24-well plates, and synchronized HEV infection was performed as described above. At 56 h and day 7 post-internalization, cells were trypsinized for 10 min, gently resuspended, and singularized by pipetting thrice through a 70 µm cell strainer. Two wells of each sample were combined. Single-cell suspensions were washed once with 0.04% BSA in PBS and finally resuspended in 0.04% BSA in PBS. Cell suspension was counted and inspected for cell death using trypan blue. Single-cell suspensions were loaded onto the 10x Chromium controller according to manufacturer’s instructions of the Chromium Next GEM Single Cell 3ʹ Kit v3.1 (10x Genomics) with a targeted cell recovery of 5,000. Sequencing libraries were prepared according to the manufacturer’s instructions. Briefly, GEMs were generated, reverse transcription was performed, GEMs were cleaned up, and cDNA was amplified and cleaned up with SPRIselect beads (Beckman Coulter). Quantification and quality control were performed using Qubit (Thermo Fisher Scientific) and the 5200 Fragment Analyzer System (Agilent Technologies). Fragmentation, end repair, and A-tailing were followed by SPRIselect-based cleanup, adaptor ligation, sample indexing, and again, SPRIselect cleanup. Quantification and quality control were performed with 5200 Fragment Analyzer System (Agilent Technologies) and NEBNext Library Quant Kit for Illumina (New England Biolabs). Resulting libraries were pooled and sequenced on Illumina NextSeq550 (high-output mode, paired-end, 150 cycles).

### scRNA-seq data analysis

Fastq files were aligned and counted using the Cell Ranger 7.1.0 pipeline. As a human genome reference we used GRCh38-3.0.0. For the gene set enrichment analysis, up- and down- regulated genes were identified using a Wilcoxon rank sum test as implemented in the FindMarkers function from Seurat 5.0. We filtered out genes which were expressed in less than 5% of cells in either uninfected or infected samples and with p-adjusted values < 0.05. GSEA was performed using the enrichR v3.2 R package. As a reference we used the hallmark gene sets of the Human MSigDB Collections^38^. The entire analysis was carried out using R 4.3.3. For clustering and UMAP projections, we used only a subset of genes based on a list of ISGs reported by Schoggins *et al.*^39^. Counts were scaled and log1-normalized. The UMAP projections were calculated over the first 20 dimensions of the PCA, setting the following parameters: min.dist=0.5 and n.neighbors=200. In order to identify clusters associated with transcriptionally active/inactive states, we set a resolution of 0.1 in the FindClusters function and then we inspected the expression of ISGs in the identified clusters. For the DEA we used the FindMarkers function and filtered out genes expressed in less than 5% of cells in the compared conditions and with p-adjusted values < 0.05. For ISG expression we used AUC values to score cells based on the Schoggins signature mentioned above implemented with the AUCell 1.24.0 R package. For the comparison of the gene expression across conditions we used a two-sided Wilcoxon rank sum test. Binarized ORF2 expression was carried out by defining positive cells as those expressing at least one HEV RNA count. We used the Seurat 5.0 R package for the entire analysis. The data is accessible under the GEO accession number GSE288400.

### Statistical analysis

Graphs were made and statistical analysis was performed using GraphPad Prism 8 or R 4.3.3 for the scRNA-seq analysis. Statistical tests and p-values are listed in the respective figure legends.

## Supporting information

Supplementary Information Mehnert et al.

## Acknowledgments

The authors gratefully acknowledge Dr. Suzanne Emerson, Dr. Rainer Ulrich, Dr. Frauke Mücksch, and Dr. Henrik Kaessmann for sharing reagents, and Andrew Freistaedter and Céline Schneider for excellent technical support. We acknowledge Dr. Vibor Laketa, head of the Infectious Diseases Imaging Platform (IDIP) at the University Hospital Heidelberg for expert support. The authors thank Dr. Marlène Dreux for critical reading of the manuscript. We thank the BIOI2 platform and the I2BC’s IT support team for making AlphaFold 2.3 easily accessible at the I2BC.

## Funding

A.M., S.S., C.R., C.H., M.B., and V.L.D.T were supported by grants from the Deutsche Forschungsgemeinschaft (DFG, German Research Foundation) – Projektnummer – 272983813 SFB/TRR 179. V.L.D.T. was additionally supported by the Chica and Heinz Schaller Foundation, DFG grant DA1640/3-1, and TTU Hepatitis Project 05.823. J.H. was supported by a fellowship from the China Scholarship Council.

## Author contributions

Conceptualization, A.M., and V.L.D.T.; Methodology, A.M., S.S., C.R, V.G.M, C.S., A.L.C., J.H., and T.T.; Investigation, A.M., S.S, C.R., and T.T.; Resources: D.K., M.K., X.W.; Software, A.M., S.S., C.R., A.L.C., T.T.; Data analysis, A.M., S.S., C.R., A.L.C., C.H., J.H.; Writing original draft, A.M., and V.L.D.T.; Final draft, A.M., and V.L.D.T.; Supervision, C.H., B.M., V.L.D.T.; Funding, V.L.D.T.

## Declaration of interests

All authors declare no conflict of interest

## Notes

### Competing Interest Statement

The authors have declared no competing interest.

### Summary of Updates

New data has been added to the manuscript.

