## Supplementary Information Mehnert et al. for "The hepatitis E virus capsid protein ORF2 counteracts cell-intrinsic antiviral responses to enable persistence in hepatocytes"

### Supplementary Methods

*Quantitative (real-time) reverse transcription PCR (RT-qPCR)*

RNA was extracted from cell lysates either with the Universal RNA kit (Roboklon), Monarch Total RNA Miniprep Kit (New England Biolabs), or with TRIzol reagent (Invitrogen), following respective manufacturer's instructions. cDNA was synthesized using the iScript cDNA Synthesis Kit (Bio-Rad) or the High Capacity cDNA Reverse Transcription Kit (Thermo Fisher Scientific) and diluted. qPCR was then performed with iTaq Universal SYBR Green Supermix (Bio-Rad) on a CFX96 Real-Time PCR Detection System (Bio-Rad) using the primers listed in Suppl. table 1. Absolute HEV genome copies were calculated from an HEV standard curve, produced by serial 10-fold dilutions of a 10 ng/µL-concentrated pBSK-HEV-p6 plasmid. *IFNL1*, *ISG15*, and *IFIT1* expression in HepG2/C3A cells, HLCs, and Huh7.5 cells was normalized over the housekeeping gene *RPS11* using the $2^{-\Delta Ct}$ method or additionally normalized to HEV RNA using the $2^{-\Delta\Delta Ct}$ method. For *IFNB1*, *TNFAIP3*, and *IFIT1* expression in A549-derived cells, relative expression over the housekeeping gene *GAPDH* was calculated using the $2^{-\Delta Ct}$ method.

*Co-immunoprecipitation (co-IP)*

$1.2 \times 10^7$ HEK293T cells were seeded on 10-cm cell culture dishes coated with 100 µg/mL poly-L-lysine. The next day, cells were co-transfected with 10 µg of plasmid encoding ORF2-HA or HA-tagged ORF2 mutants and 10 µg of plasmid encoding TBK1-V5 in cDMEM without pen/strep. Polyethyleneimine (PEI) was used for transfection at a DNA:PEI ratio of 1:3. Medium was changed after 5-6 h. After 24 h, cells were washed twice with cold PBS and harvested by scraping in cold PBS containing 0.5x cOmplete Mini Protease Inhibitor Cocktail (Roche). Cells were pelleted and resuspended in 1 mL cold lysis buffer (25 mM Tris-HCl, pH 7.5, 150 mM NaCl, 1 mM EDTA, 1% NP-40, 5% glycerol) containing 1x cOmplete Mini Protease Inhibitor Cocktail (Roche) and incubated on ice for 30 minutes. After removal of DNA, 6% of the lysate was set aside as input and boiled with Laemmli sodium dodecyl sulfate (SDS) sample buffer at 95 °C for 10 min. 20 µL Pierce Anti-HA Magnetic Beads (Thermo Fisher Scientific) were added to each sample and incubated under rotation for 2 h at 4 °C. Beads were washed thrice with lysis buffer containing 1x cOmplete Mini Protease Inhibitor Cocktail (Roche) and eluted in Laemmli SDS sample buffer by boiling at 95 °C for 10 min.

*RNA fluorescence* in situ *hybridization and quantification*

Fixed, electroporated HepG2/C3A cells were stained using the RNAscope Multiplex Fluorescent V2 Assay (ACDBio), following the manufacturer's protocol, starting from the incubation step with hydrogen peroxide. HEV RNA was specifically targeted with the V-HEV-p6-ORF2 probe (ACDBio, cat no. 586651), detecting both genomic and subgenomic HEV RNA. Cells were counterstained with Hoechst (Thermo Fisher Scientific) and imaged using an inverted Nikon Eclipse Ts2-FL widefield fluorescence microscope. For quantification, HEV RNA signal was segmented in ilastik[1] and then overlaid with nuclei, which were segmented in CellProfiler[2]. The percentage of HEV RNA-positive cells was obtained using CellProfiler. For representation, images were merged and adjusted with Fiji[3].

*Western blot*

Co-IP samples were prepared as described above. Electroporated S10-3 and S10-3/ORF2 cells were lysed in Pierce RIPA buffer (Thermo Fisher Scientific) with 1x cOmplete Mini Protease Inhibitor Cocktail (Roche) on ice for 30 min, supplemented with Laemmli SDS sample buffer, boiled at 95 °C for 10 min. For detection of ORF3, samples were immediately run on SDS-polyacrylamide gel electrophoresis (PAGE) gels. Proteins were transferred and membranes were blocked with 5% milk (Carl Roth) in PBS/0.1% Tween-20 (PBS-T). The following antibodies were incubated overnight in 5% milk/PBS-T at 4 °C: mouse-anti-β-actin, 1:4000 (Sigma-Aldrich, cat no. A2228); mouse-anti-ORF2 1E6, 1:500 (Merck, cat no. MAB8002); mouse-anti-ORF3, 1:25 (University of Geneva Antibody Facility, cat no. ABCD_RB198); rabbit-anti-HA, 1:1000 (Cell Signaling Technology, cat. no. 3724); rabbit-anti-V5, 1:1000 (Cell Signaling Technology, cat. no. 13202). After three washes with PBS-T, membranes were incubated with horseradish peroxidase (HRP)-coupled anti-mouse or anti-rabbit secondary antibodies (Jackson Immunoresearch, 1:4000) for 1 h at room temperature. Membranes were washed thrice with PBS-T and once with PBS, and chemiluminescent signal was developed using Pierce ECL Western Blotting Substrate (Thermo Fisher Scientific).

*Generation of human pluripotent stem cell (hPSC)-derived hepatocyte-like cells (HLCs) and HEV infection*

The hESC cell line RUES2[4] was cultured in mTeSR1 (STEMCELL Technologies) on cell culture plates coated with Matrigel (Corning). RUES2 cells were differentiated to definitive endoderm (DE) using the STEMdiff Definitive Endoderm Differentiation Kit (STEMCELL Technologies) according to manufacturer's instructions. As described previously[5], DE cells were reseeded and differentiated to hepatocyte progenitors for five days in basal Rosewell Park Memorial Institute (RPMI) 1640 medium with HEPES (Gibco), supplemented with B-27 custom supplement (Gibco), GlutaMAX (Gibco), NEAA (Gibco), and pen/strep (Gibco), additionally containing the human growth factors bone morphogenetic protein 4 (BMP4, PeproTech) and fibroblast growth factor basic (FGFb, Gibco). Immature hepatocytes were obtained after five days in basal RPMI containing human hepatocyte growth factor (HGF, PeproTech). Maturation into hepatocytes was achieved by final differentiation in the supplemented Hepatocyte Culture Medium BulletKit (HCM, Lonza; no HEGF component) containing human oncostatin M (OSM, R&D Systems). Mature HLCs were infected with equal genome copies of HEV WT and ΔORF2 (30 GE/cell) in HCM/OSM medium. The next day, the inoculum was removed and cells were washed twice with Dulbecco's Balanced Salt Solution (DPBS, Gibco). HCM/OSM medium was replenished every two days. Cells were fixed with 4% PFA or lysed for RNA extraction with TRIzol at respective time points.

*Immunofluorescence (IF) staining*

Samples were fixed with 4% PFA and permeabilized/blocked in 10% goat serum (MP Biomedicals), 1% bovine serum albumin (BSA, Carl Roth), and 0.1% Triton X-100 in PBS. Primary antibodies were incubated overnight at 4 °C in blocking/permeabilization solution with the following dilutions: rabbit-anti-ORF2, 1:6000 (a kind gift from Rainer Ulrich[6]); mouse-anti-albumin, 1:1000 (tebubio, cat. no. CL2513A). Samples were washed thrice with PBS and incubated with Alexa Fluor-conjugated secondary antibodies (1:1000, Thermo Fisher Scientific) for 1 h at room temperature. After washing thrice, cells were counterstained with Hoechst (1:1000, Thermo Fisher Scientific) and imaged using an inverted Nikon Eclipse Ts2-FL widefield fluorescence microscope. Images were analyzed and merged with Fiji[3].

*IFNβ ELISA*

Supernatants from A549 cells challenged with Mengo-Zn virus, Sendai virus (SeV), and poly(I:C) supernatant feeding for 24 h were collected and IFNβ secretion was measured using a bioluminescent human IFNβ ELISA (LumiKine™ Xpress hIFN-β 2.0, Invivogen) according to manufacturer's instructions.

*Electroporation of A549 cells*

$2x10^6$ cells were pelleted and resuspended in 200 µL cytomix (120 mM KCl, 0.15 mM $CaCl_2$, 10 mM $KPO_4$, 25 mM HEPES, 2 mM EGTA, and 2 mM $MgCl_2$), and transferred to a 0.2 cm cuvette containing 500 ng high molecular weight (HMW) poly(I:C) (Invivogen) or control poly(C) (Sigma-Aldrich). Electro-transfection was performed at 166 V and 500 µF using the Gene Pulser Xcell modular electro-transfection system (Bio-Rad). Transfected cell suspensions were transferred to pre-warmed DMEM, centrifuged, and resuspended in 4 mL cDMEM plus 1% NEAA. 500 µL of the final cell suspension were seeded in 24-well plates.

*Cell viability (MTS) assay*

After EPO of HepG2/C3A with HEV WT RNA or respective mutants, cells were mixed 1:1 with mock-electroporated cells and $3x10^4$ cells were seeded in triplicates on 96-well plates. On day 1, day 3, and day 5 post-EPO, cDMEM was replaced and 6 µM TBK1 inhibitor BX795 (InvivoGen) or corresponding DMSO vehicle control was added to the wells dedicated to analysis 48 h later. At the respective time points, CellTiter 96 AQueous One Solution Cell Proliferation Assay (Promega) was performed according to manufacturer's instructions. Cells were incubated for 1 h and absorbance at 490 nm was measured using a plate reader (Tecan).

*Computational prediction of ORF2 and TBK1 interaction using AlphaFold 2.3*

To predict interactions between ORF2 and TKB1, we constructed 13 distinct systems, varying the chunking strategy and oligomerization states of both proteins. For each system, five structural models were generated using three different random seeds, resulting in a total of 195 models. These models were produced using an in-house implementation of ColabFold[7], employing AlphaFold-Multimer 2.3 weights[8,9] and a multiple sequence alignment computed with the MMSEQS2 webserver[10-13]. Model quality was assessed using pLDDT[14], Predicted Aligned Error, and actifPTM (ACTual InterFace PTM) scores[15]. Additionally, we computed pdock, pdock2, and LIS scores for each model using the AF_analysis package (https://github.com/samuelmurail/af_analysis). Contact maps were generated in-house using the MDTraj Python library[16]. A contact between two amino acids was defined as the presence of at least two atoms within a distance of 5 Å. All scripts used and models produced in this study are available on Zenodo (https://doi.org/10.5281/zenodo.14751497).

### Supplementary Figures

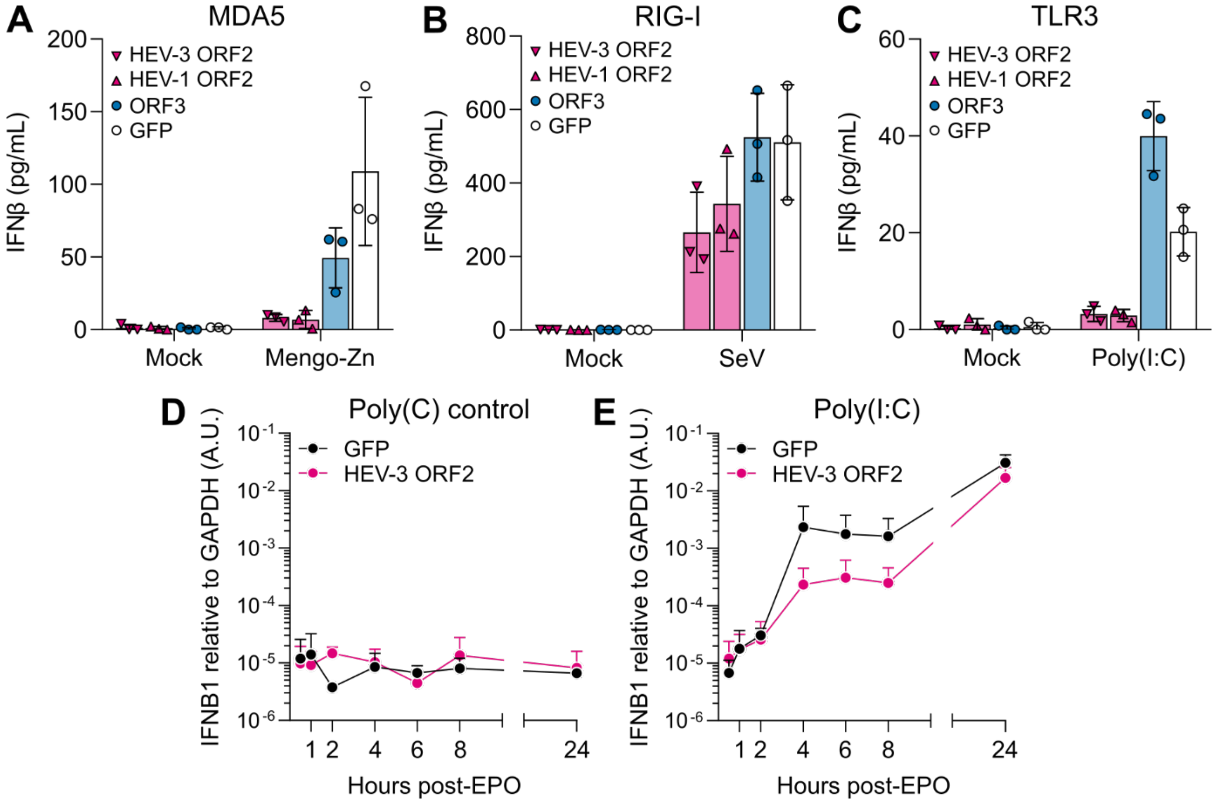

**Supplementary figure 1: The HEV ORF2 protein interferes with the secretion of IFNβ but does not affect the expression kinetics.**

(A) A549 cells harboring knockouts of the PRRs RIG-I and MDA5 and ectopically expressing a single PRR (MDA5, RIG-I, or TLR3) together with either HEV-3 ORF2, HEV-1 ORF2, ORF3, or GFP were challenged with either Mengo-Zn virus at MOI 1, (B) Sendai virus (SeV) at MOI 0.75, or (C) 50 µg/mL poly(I:C) supernatant feeding for 24 h. Supernatant was collected and IFNβ was measured by enzyme-linked immunosorbent assay (ELISA). (D) A549 cells harboring knockouts of the PRRs RIG-I and MDA5 and ectopically expressing MDA5 and either GFP or HEV-3 ORF2 were electroporated with poly(C) control or (E) poly(I:C). At indicated time points post-EPO, *IFNB1* expression was analyzed by RT-qPCR relative to the housekeeping gene *GAPDH* using the $2^{-\Delta Ct}$ method. Data shown mean ± SD of n = 3 independent biological experiments.

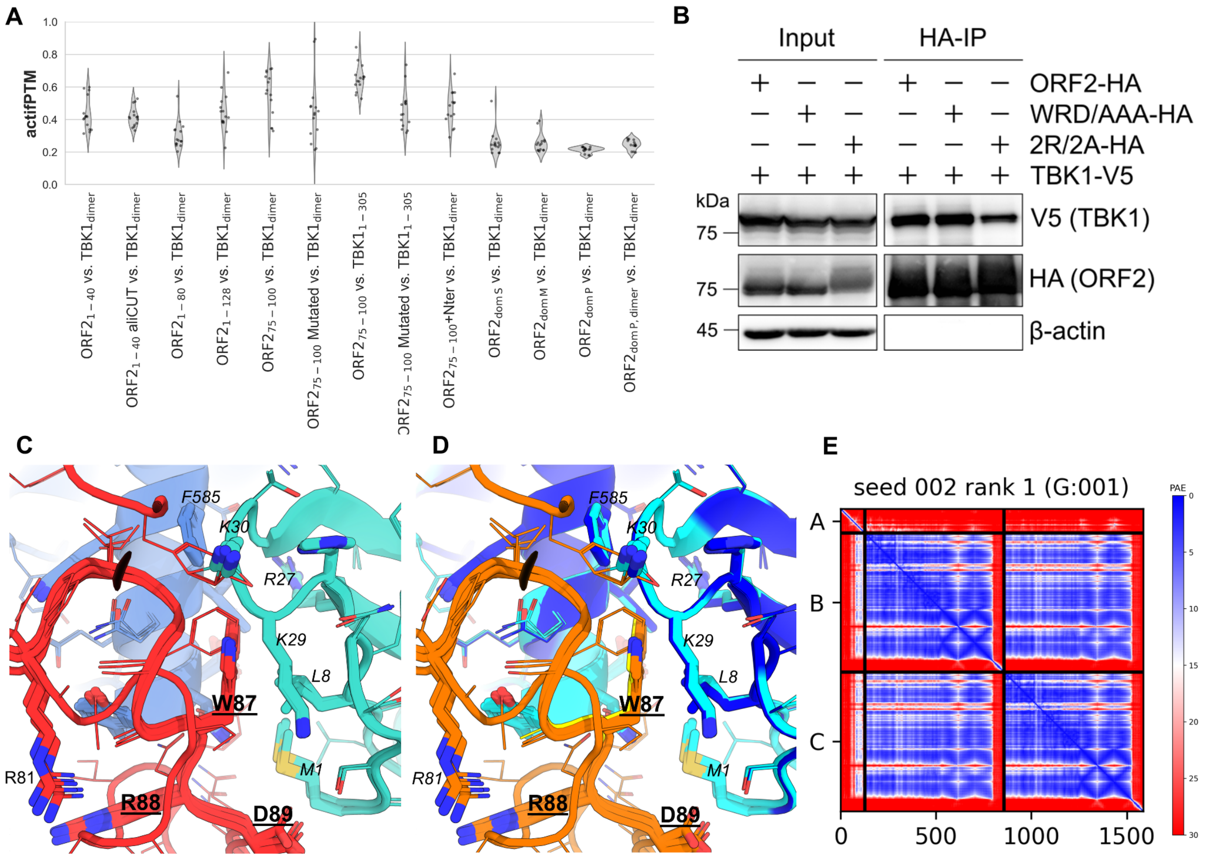

**Supplementary figure 2: AlphaFold 2.3 modeling of the ORF2-TBK1 interaction.**

(A) actifPTM scores assessing predicted interactions between segments of ORF2 and TBK1. Higher scores indicate stronger predicted interactions. (B) HEK293T cells were transfected with ORF2-HA, ORF2-WRD/AAA-HA, or ORF2-2R/2A-HA and V5-tagged TBK1 and lysed 24 h post-transfection. Anti-HA co-IP and WB analysis for TBK1 (anti-V5 staining), ORF2 (anti-HA staining), and β-actin were performed. Representative blot of n = 2 independent biological experiments. (C-D) Structural analysis of the top five ranked models for the interaction between ORF2 residues 1-128 (red) and a TBK1 dimer. Panel C highlights the molecular environment surrounding ORF2 W87, color-coded by domain (TBK1 kinase domain in turquoise, C-terminal domain in blue). Panel D depicts the same region colored according to pLDDT scores, reflecting model confidence (orange – very low; yellow – low; cyan – high; blue – very high). Modeling convergence of the models and the chemical environment of W87 and R88 suggest potential for stabilizing cation-π interactions. (E) Predicted Aligned Error (PAE) for the highest-ranking ORF21-128 vs. TBK1 dimer model. Low PAE values in the N-terminal region of ORF21-128, particularly with respect to TBK1 chains B and C, indicate a likely accurate placement of this region relative to the TBK1 monomers.

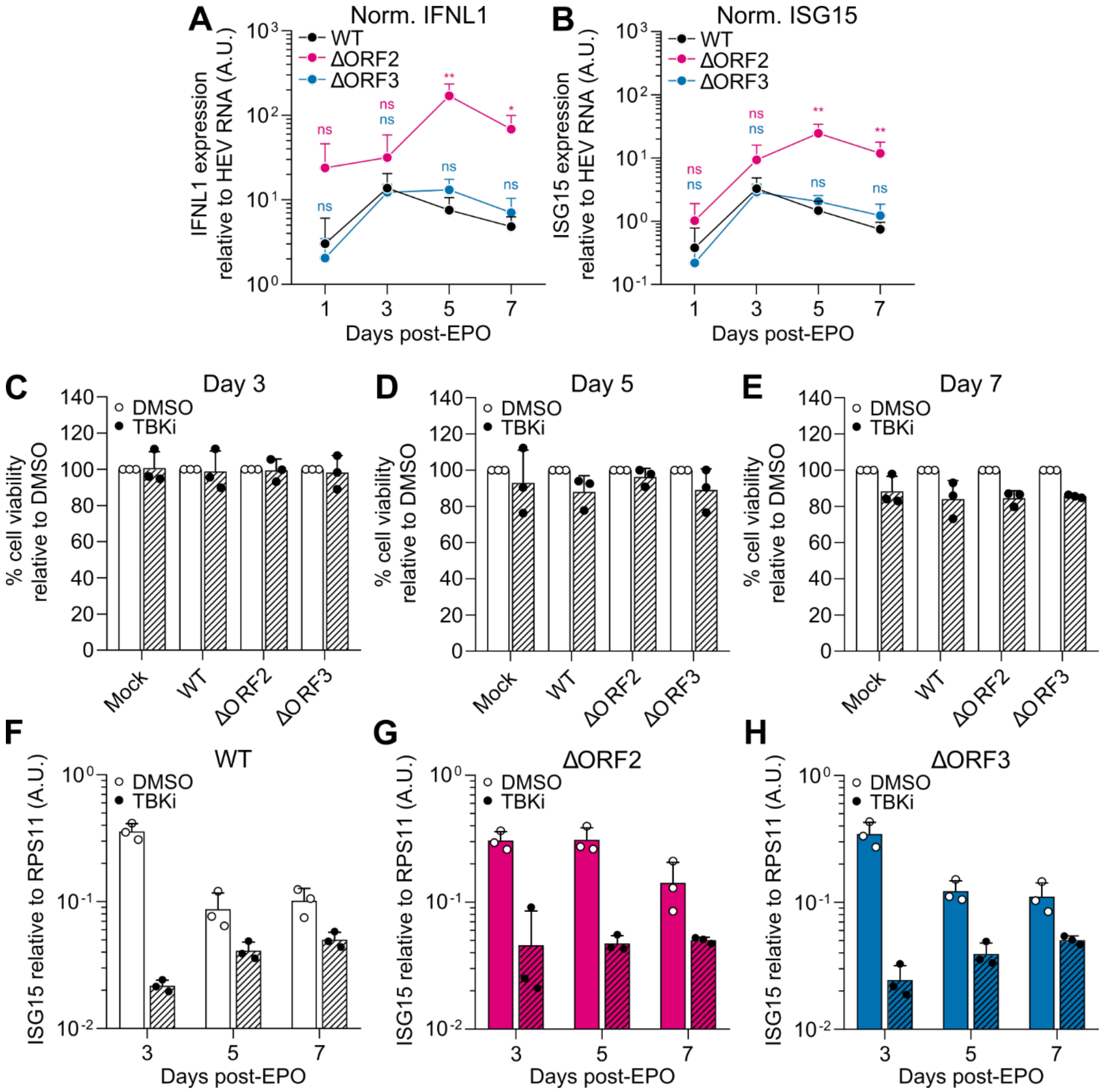

**Supplementary figure 3: Normalized *IFNL1* and *ISG15* expression over HEV RNA and effect of TBK1 inhibitor BX795 on cell viability and *ISG15* expression.**

(A-B) Normalized data of Fig. 2B-D over HEV RNA: Expression of (A) *IFNL1* and (B) *ISG15* upon EPO of HepG2/C3A cells with HEV WT, ΔORF2, or ΔORF3 were determined relative to the housekeeping gene *RPS11* and normalized to HEV RNA using the $2^{-\Delta\Delta Ct}$ method. Data show mean ± SD of n = 3 independent biological experiments. Statistical analysis was performed using one-way ANOVA of each time point independently and comparison to WT is indicated above the respective time points in the corresponding colors. *: $p < 0.05$; **: $p < 0.01$; ns, non-significant. A.U., arbitrary units; norm., normalized. (C) HepG2/C3A cells were either mock-electroporated or electroporated with HEV WT, ΔORF2, ΔORF3 RNA and additionally treated with 6 µM of the TBK1 inhibitor BX795 (TBKi) or respective DMSO vehicle control 48 h prior to the time point of harvest. Cell viability was determined at day 3, (D) day 5, and (E) day 7 post-EPO using the CellTiter 96® $AQ_{ueous}$ One Solution Cell Proliferation Assay (Promega). Data was normalized to the respective DMSO controls. Data show mean ± SD of n = 3 independent biological experiments. (F) HepG2/C3A cells electroporated with WT, (G) ΔORF2, or (H) ΔORF3 RNA and treated with 6 µM TBKi or respective vehicle control were analyzed for *ISG15* expression relative to the housekeeping gene *RPS11* by RT-qPR at indicated time points post-EPO using the $2^{-\Delta Ct}$ method. Data show mean ± SD of n = 3 independent biological experiments.

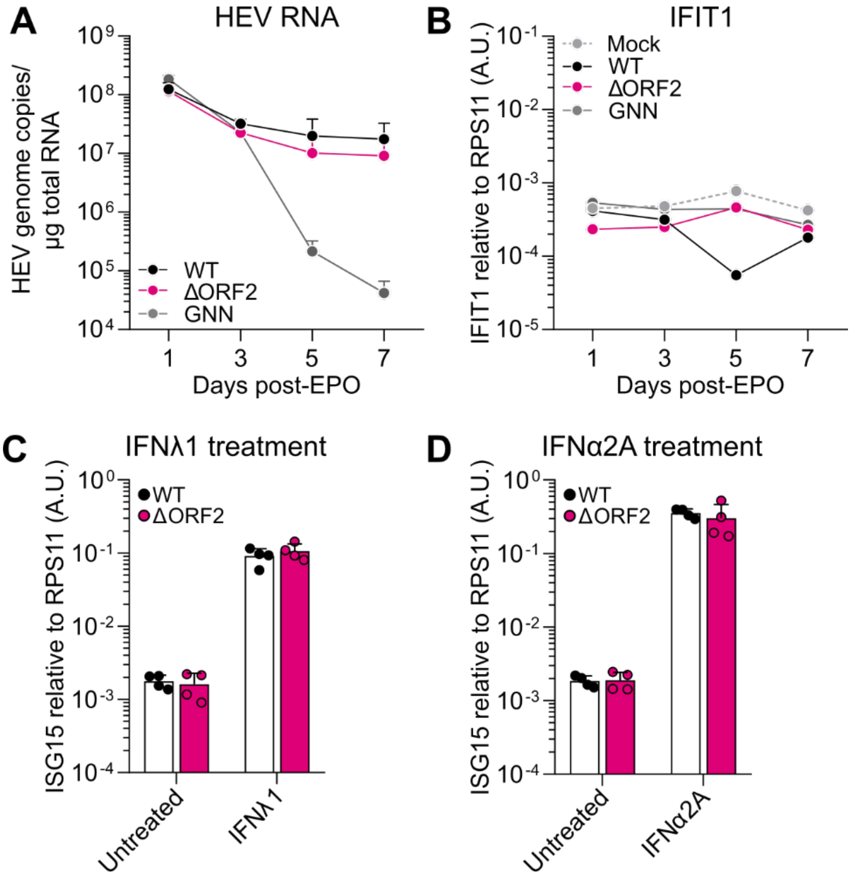

**Supplementary figure 4: HEV WT and ΔORF2 replicate to similar levels in immunodeficient Huh7.5 cells and exogenous IFN stimulation induces ISG expression.**

(A) Huh7.5 cells were electroporated with *in vitro* transcribed HEV WT, ΔORF2, or replication-incompetent GNN RNA and analyzed at indicated time points post-electroporation (EPO) for HEV RNA by RT-qPCR. Data show mean ± SEM of n = 2 independent biological experiments. (B) Electroporated Huh7.5 cells from (A) were analyzed by RT-qPCR for expression of the ISG *IFIT1* relative to the housekeeping gene RPS11 using the $2^{-\Delta Ct}$ method. Data shows a single biological experiment. (C) Huh7.5 cells electroporated with HEV WT or ΔORF2 RNA and treated with 10 ng/mL IFNλ1 or (D) 10,000 IU/mL IFNα2A from day 4 to day 7 post-EPO were analyzed for *ISG15* expression at day 7 post-EPO by RT-qPCR. *ISG15* expression was analyzed using the $2^{-\Delta Ct}$ method. Data show mean ± SD of n = 4 biological repeats from two independent biological experiments.

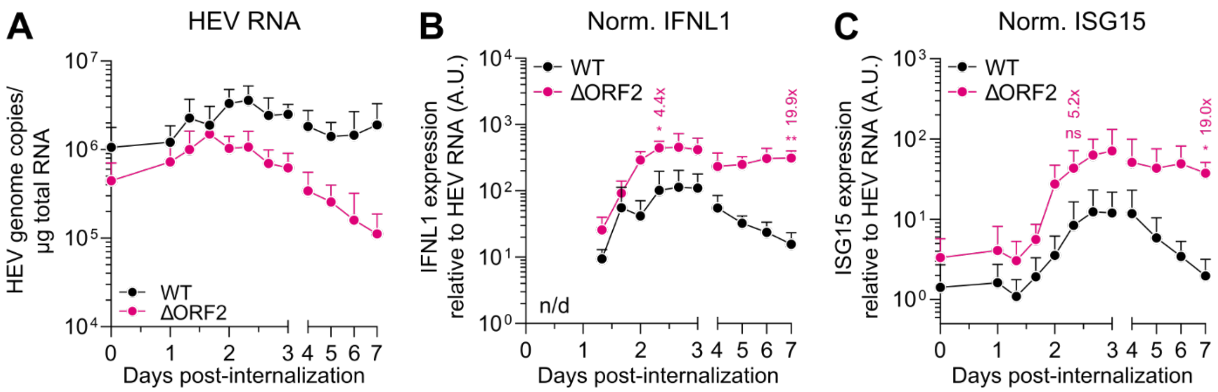

**Supplementary figure 5: Non-normalized HEV RNA and normalized antiviral response gene expression over HEV RNA of HEV WT and ΔORF2 infection.**

(A) Non-normalized data of Figure 3C: Equal genome copies of HEV WT and ΔORF2 virus particles (30 GE/cell) were bound on HepG2/C3A cells for 2 h at 4 °C prior to internalization at 37 °C for 8 h, followed by removal of inoculum (= day 0). RT-qPCR was performed at indicated time points post-internalization to determine HEV genome copies. (B) Normalized data of Figure 3D-E over HEV RNA: *IFNL1* expression relative to the housekeeping gene RPS11, and (C) *ISG15* expression relative to *RPS11* were additionally normalized over HEV RNA using the $2^{-\Delta\Delta Ct}$ method. Statistical analysis of fold changes of ΔORF2 over WT are indicated above the respective time points in the corresponding color. Data show mean ± SD of n = 3 independent biological experiments. Statistical analysis was performed using unpaired two-tailed Student's t-test of the respective days independently. *: $p < 0.05$; **: $p < 0.01$; ns, non-significant. A.U., arbitrary units; n/d, not detectable; norm., normalized.

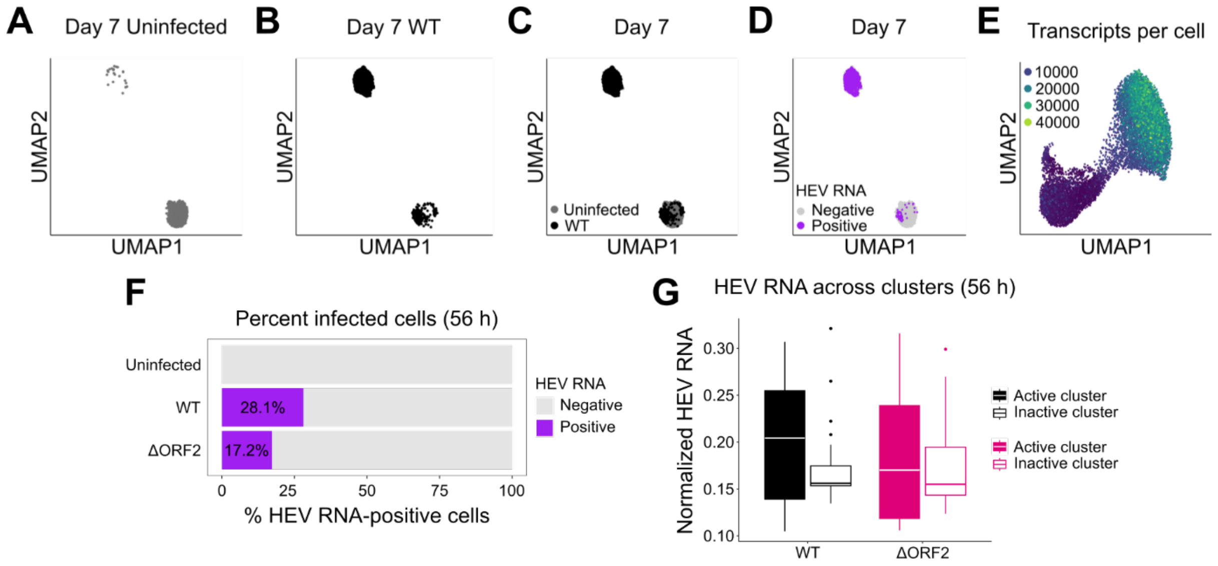

**Supplementary figure 6: scRNA-seq analysis of WT infection at day 7 and analysis of infected cells across clusters at 56 h.**

(A) UMAP projections of uninfected, (B) WT-infected, and (C) combined UMAP projections of uninfected and WT-infected HepG2/C3A cells harvested at day 7 post-infection for scRNA-seq analysis, colored by condition. Cells were clustered based on a list of approximately 400 ISGs published previously by Schoggins *et al.*[17]. (D) Combined UMAP projection of the uninfected and WT-infected samples at day 7 post-infection with indicated binarized HEV RNA counts in purple. (E) UMAP projection of uninfected, WT-, and ΔORF2-infected samples at 56 h post-infection, highlighting the number of transcripts detected per cell in all samples. (F) The percentage of infected cells, defined by at least one HEV RNA count detected per cell, was quantified in uninfected, WT-, and ΔORF2-infected samples at 56 h post-infection. (G) Normalized HEV RNA counts across infected cells in the inactive and active clusters of WT- and ΔORF2-infected samples at 56 h post-infection.

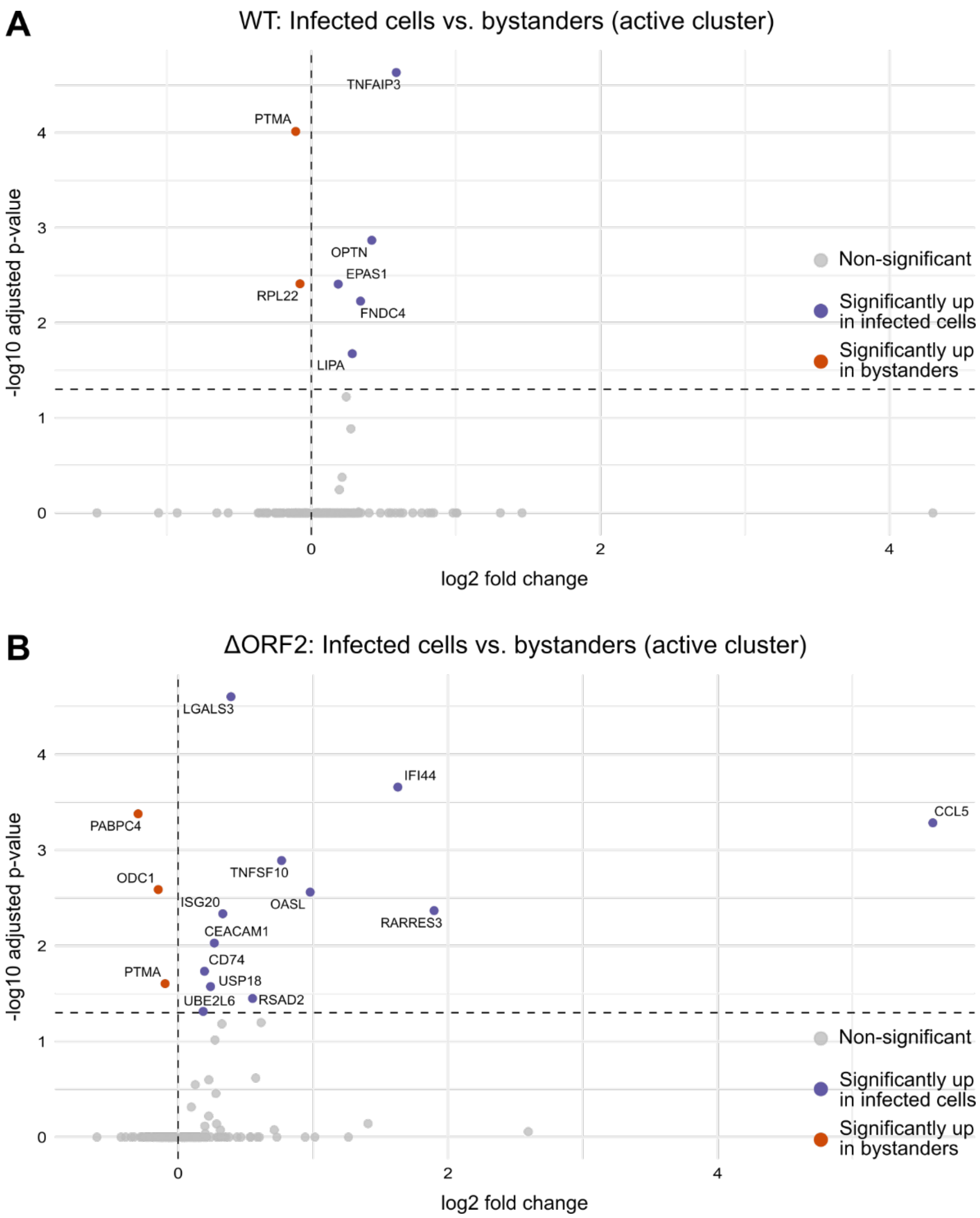

**Supplementary figure 7: Specific ISG signatures of infected cells and bystanders in WT and ΔORF2 infection.**

(A) ISG expression according to the list of approximately 400 ISGs published previously by Schoggins *et al.*[17] was compared at 56 h post-synchronized infection in WT- and (B) ΔORF2-infected HepG2/C3A cells between infected cells (defined by detection of at least one HEV RNA copy per cell) and bystanders in the active cluster using a Wilcoxon rank sum test. Significant genes with a log2 fold change > 0 were specifically upregulated in infected cells while significant genes with a log2 fold change < 0 were specifically upregulated in bystanders.

**Supplementary table 1: List of oligonucleotides**

| Oligonucleotide | Target | Sequence (5'-3') | Ref |
|---|---|---|---|
| HEV fw | HEV | GGTGGTTTCTGGGGTGAC | Wu *et al.*[5] |
| HEV rv | | AGGGGTTGGTTGGATGAA | Wu *et al.*[5] |
| RPS11 fw | RPS11 | GCCGAGACTATCTGCACTAC | Wu *et al.*[5] |
| RPS11 rv | | ATGTCCAGCCTCAGAACTTC | Wu *et al.*[5] |
| IFNL1 fw | IFNL1 | GTGACTTTGGTGCTAGGCTTG | Wu *et al.*[5] |
| IFNL1 rv | | GCCTCAGGTCCCAATTCCC | Wu *et al.*[5] |
| ISG15 fw | ISG15 | CTGTTCTGGCTGACCTTCG | Wu *et al.*[5] |
| ISG15 rv | | GGCTTGAGGCCGTACTCC | Wu *et al.*[5] |
| GAPDH fw | GAPDH | TCGGAGTCAACGGATTTGGT | Wüst *et al.*[18] |
| GAPDH rv | | TTCCCGTTCTCAGCCTTGAC | Wüst *et al.*[18] |
| IFNB1 fw | IFNB1 | CGCCGCATTGACCATCTA | Wüst *et al.*[18] |
| IFNB1 rv | | GACATTAGCCAGGAGGTTCTC | Wüst *et al.*[18] |
| TNFAIP3 (A20) fw | TNFAIP3 | TCCTCAGGCTTTGTATTTGAGC | Wüst *et al.*[18] |
| TNFAIP3 (A20) rv | | TGTGTATCGGTGCATGGTTTTA | Wüst *et al.*[18] |
| IFIT1 fw | IFIT1 | GAATAGCCAGATCTCAGAGGAGC | Wüst *et al.*[18] |
| IFIT1 rv | | CCATTTGTACTCATGGTTGCTGT | Wüst *et al.*[18] |
| ΔORF2 mutant first start codon fw | HEV ORF2 | TGGGATCACCGTGTGCCCTAG | This study |
| ΔORF2 mutant first start codon rv | | CTAGGGCACACGGTGATCCCA | This study |
| ΔORF2 mutant second start codon fw | HEV ORF2 | GTTTCTGCCTGTGCTGCCCG | This study |
| ΔORF2 mutant second start codon rv | | CGGGCAGCACAGGCAGAAAC | This study |
| ΔORF3 mutant fw | HEV ORF3 | CATCGCCCAGCGGATCACCAT | This study |
| ΔORF3 mutant rv | | ATGGTGATCCGCTGGGCGATG | This study |
| GNN mutant fw | HEV ORF1 | GCCTTTAAGGGTAATAATTCGGTGGT | This study |
| GNN mutant rv | | ACCACCGAATTATTACCCTTAAAGGC | This study |
| ORF2 2R/2A mutant fw | HEV ORF2 | GTCGTCGTGGGGCGGCCAGCGGCGGTG | Hervouet *et al.*[19] |
| ORF2 2R/2A mutant rv | | GCACCGCCGCTGGCCGCCCCACGACGACG | Hervouet *et al.*[19] |
| ORF2 WRD/AAA mutant fw | HEV ORF2 | CTTGGCTCCGCTGCGGCTGCCCAGTCCCAG | This study |
| ORF2 WRD/AAA mutant rv | HEV ORF2 | CTGGGACTGGGCAGCCGCAGCGGAGCCAAG | This study |

**Supplementary table 2: Intersection list of ISGs for Venn diagram in Fig. 4L**

| Intersection | WT only | ΔORF2 only |
|---|---|---|
| ABTB2 | AKT3 | APOL1 |
| ACSL1 | ALDH1A1 | APOL2 |
| ADAR | ATF3 | BCL3 |
| ADM | C1S | BLZF1 |
| ANGPTL1 | CD74 | CD274 |
| ANKFY1 | FUT4 | CFB |
| APOBEC3G | GTPBP1 | CREB3L3 |
| APOL3 | HES4 | DCP1A |
| APOL6 | IRF1 | ENPP1 |
| ARG2 | LIPA | ETV6 |
| ARHGEF3 | MOV10 | EXT1 |
| ARNTL | MT1G | FNDC3B |
| ATP10D | NPAS2 | GBP3 |
| B2M | ODC1 | HLA-E |
| B4GALT5 | PFKFB3 | IFI44 |
| BAG1 | SECTM1 | IFI44L |
| BATF2 | SLC1A1 | IFNGR1 |
| BCL2L14 | | JAK2 |
| BLVRA | | LAMP3 |
| BST2 | | LGMN |
| BTN3A3 | | MAFF |
| BUB1 | | MYD88 |
| C4orf33 | | NFIL3 |
| CCDC92 | | OPTN |
| CCND3 | | RARRES3 |
| CD9 | | RBCK1 |
| CDKN1A | | RNF24 |
| CEACAM1 | | RTP4 |
| CEBPD | | SCARB2 |
| CHMP5 | | SLC25A30 |
| CNP | | SPSB1 |
| COMMD3 | | TMEM140 |
| CPT1A | | |
| CRY1 | | |
| CX3CL1 | | |
| DDX3X | | |
| DDX58 | | |
| DDX60 | | |
| DEFB1 | | |
| DHX58 | | |
| DTX3L | | |
| EHD4 | | |
| EIF2AK2 | | |

| |
|---|
| EIF3L |
| ELF1 |
| EPAS1 |
| ERLIN1 |
| ETV7 |
| FKBP5 |
| GAK |
| GALNT2 |
| GBP1 |
| GBP2 |
| GCA |
| GCH1 |
| GK |
| GLRX |
| HEG1 |
| HERC6 |
| HESX1 |
| HK2 |
| HLA-C |
| HLA-G |
| HSH2D |
| IFI27 |
| IFI30 |
| IFI35 |
| IFI6 |
| IFIH1 |
| IFIT1 |
| IFIT2 |
| IFIT3 |
| IFIT5 |
| IFITM1 |
| IFITM2 |
| IFITM3 |
| IGFBP2 |
| IL15 |
| IL15RA |
| IL17RB |
| IL1RN |
| IMPA2 |
| IRF2 |
| IRF9 |
| ISG15 |
| ISG20 |
| JUNB |
| LAP3 |
| LEPR |

| |
|---|
| LGALS3 |
| LGALS9 |
| LRG1 |
| LY6E |
| MAP3K14 |
| MARCKS |
| MASTL |
| MAX |
| MCL1 |
| MICB |
| MT1F |
| MT1X |
| MTHFD2L |
| MX1 |
| MX2 |
| N4BP1 |
| NAPA |
| NCOA3 |
| NDC80 |
| NMI |
| NUP50 |
| OAS1 |
| OAS2 |
| OAS3 |
| OASL |
| OGFR |
| P2RY6 |
| PABPC4 |
| PARP12 |
| PDGFRL |
| PDK1 |
| PHF11 |
| PIM3 |
| PLEKHA4 |
| PLSCR1 |
| PMAIP1 |
| PML |
| PMM2 |
| PNPT1 |
| PNRC1 |
| PPM1K |
| PRKD2 |
| PSMB8 |
| PSMB9 |
| PTMA |
| PUS1 |

| |
|---|
| PXK |
| RAB27A |
| RASSF4 |
| RBM25 |
| RIPK2 |
| RNASE4 |
| RNF19B |
| RPL22 |
| RSAD2 |
| SAMD4A |
| SAMHD1 |
| SAT1 |
| SERPINB9 |
| SERPINE1 |
| SLC15A3 |
| SLC16A1 |
| SLC25A28 |
| SLFN5 |
| SMAD3 |
| SNN |
| SOCS1 |
| SOCS2 |
| SP110 |
| SPTLC2 |
| SSBP3 |
| STARD5 |
| STAT1 |
| STAT2 |
| TAP1 |
| TAP2 |
| TBX3 |
| TCF7L2 |
| TDRD7 |
| TIMP1 |
| TLK2 |
| TMEM51 |
| TNFAIP3 |
| TNFRSF10A |
| TNFSF10 |
| TRAFD1 |
| TRIM14 |
| TRIM21 |
| TRIM25 |
| TRIM38 |
| TRIM5 |
| TRIM56 |

| |
|---|
| TXNIP |
| TYMP |
| UBA7 |
| UBE2L6 |
| ULK4 |
| UNC93B1 |
| UPP2 |
| USP18 |
| VAMP5 |
| WARS |
| XAF1 |
| ZNF107 |
| ZNF385B |
